# Pyramidal neurons proportionately alter the identity and survival of specific cortical interneuron subtypes

**DOI:** 10.1101/2024.07.20.604399

**Authors:** Sherry Jingjing Wu, Min Dai, Shang-Po Yang, Cai McCann, Yanjie Qiu, Vipin Kumar, Giovanni J. Marrero, Jeremiah Tsyporin, Shuhan Huang, David Shin, Jeffrey A. Stogsdill, Daniela J. Di Bella, Qing Xu, Bin Chen, Samouil L. Farhi, Evan Z. Macosko, Fei Chen, Gord Fishell

**Author notes:** Corresponding Author: Gord Fishell, Ph.D., Professor of Neurobiology and Broad Institute Member, Harvard Medical School and the Stanley Center at the Broad, Rm 201, Armenise Bldg., 220 Longwood Ave, Boston, MA 02115, USA, (HMS), (Broad). These authors contributed equally to this work (Sherry Jingjing Wu, Min Dai).

## Abstract

The mammalian cerebral cortex comprises a complex neuronal network that maintains a precise balance between excitatory pyramidal neurons and inhibitory interneurons. Accumulating evidence indicates that specific interneuron subtypes form stereotyped microcircuits with distinct pyramidal neuron classes. Here we show that pyramidal neurons play an active role in this process by promoting the survival and terminal differentiation of their associated interneuron subtypes. In wild-type cortex, interneuron subtype abundance mirrors the prevalence of their pyramidal neuron partners. In *Fezf2* mutants, which lack layer 5b pyramidal neurons and are expanded in layer 6 intratelencephalic neurons, corresponding subtype-specific shifts occur through two distinct mechanisms: somatostatin interneurons adjust their programmed cell death, whereas parvalbumin interneurons switch their subtype identity. Silencing neuronal activity or blocking vesicular release in L5b pyramidal neurons revealed that their communication with interneurons does not require voltage-gated synaptic activity and engages both tetanus toxin-sensitive and -insensitive pathways. Moreover, a targeted bioinformatic screen for ligand–receptor pairs displaying subtype-specific expression and reduced expression of pyramidal neuron-derived ligand in *Fezf2* mutants identified candidate secreted factors and adhesion molecules. These findings reveal distinct, pyramidal neuron–driven mechanisms for sculpting interneuron diversity and integrating them into local cortical circuits.

---

Single-cell sequencing has revealed a paradox of cortical organization: while interneuron subtypes are transcriptomically stable across regions^1–3^, they form specific microcircuits with regionally diverse pyramidal neuron (PN) partners^4–10^. This precision suggests that local PNs provide instructive cues for interneuron specification and integration. Here, we provide direct evidence that PNs sculpt local interneuron populations. We show that interneuron subtype composition tracks local PN populations, and that genetically re-specifying PN identity reshapes the abundance and position of corresponding somatostatin (SST) and parvalbumin (PVALB) interneurons. These changes are driven by two distinct mechanisms: PNs promote the survival of their specific SST partners while induce fate specification within the PVALB lineage. Remarkably, these PN-derived signaling are largely independent of voltage-gated synaptic activity, implicating secreted factors and cell-adhesion molecules as the key developmental cues.

## Interneuron subtypes exhibit stereotyped laminar distributions across cortical regions

To define interneuron subtypes that represent putative circuit-level components, we analyzed a P28 snRNA-seq dataset^11^, resolving 19 subtypes within the PVALB and SST interneuron classes (Fig. 1a, Extended Data Fig. 1a,b). We examined regional transcriptomic variation of individual interneuron subtypes by comparing them across the secondary motor cortex (MOs) and the primary visual cortex (VISp), two regions at opposite ends of the anterior-posterior axis. Consistent with previous work^2^, most subtypes were transcriptomically stable across regions, although our analysis revealed that SST interneurons exhibited greater regional variation than PVALB interneurons (Extended Data Fig. 1c,d).

**Figure 1.**
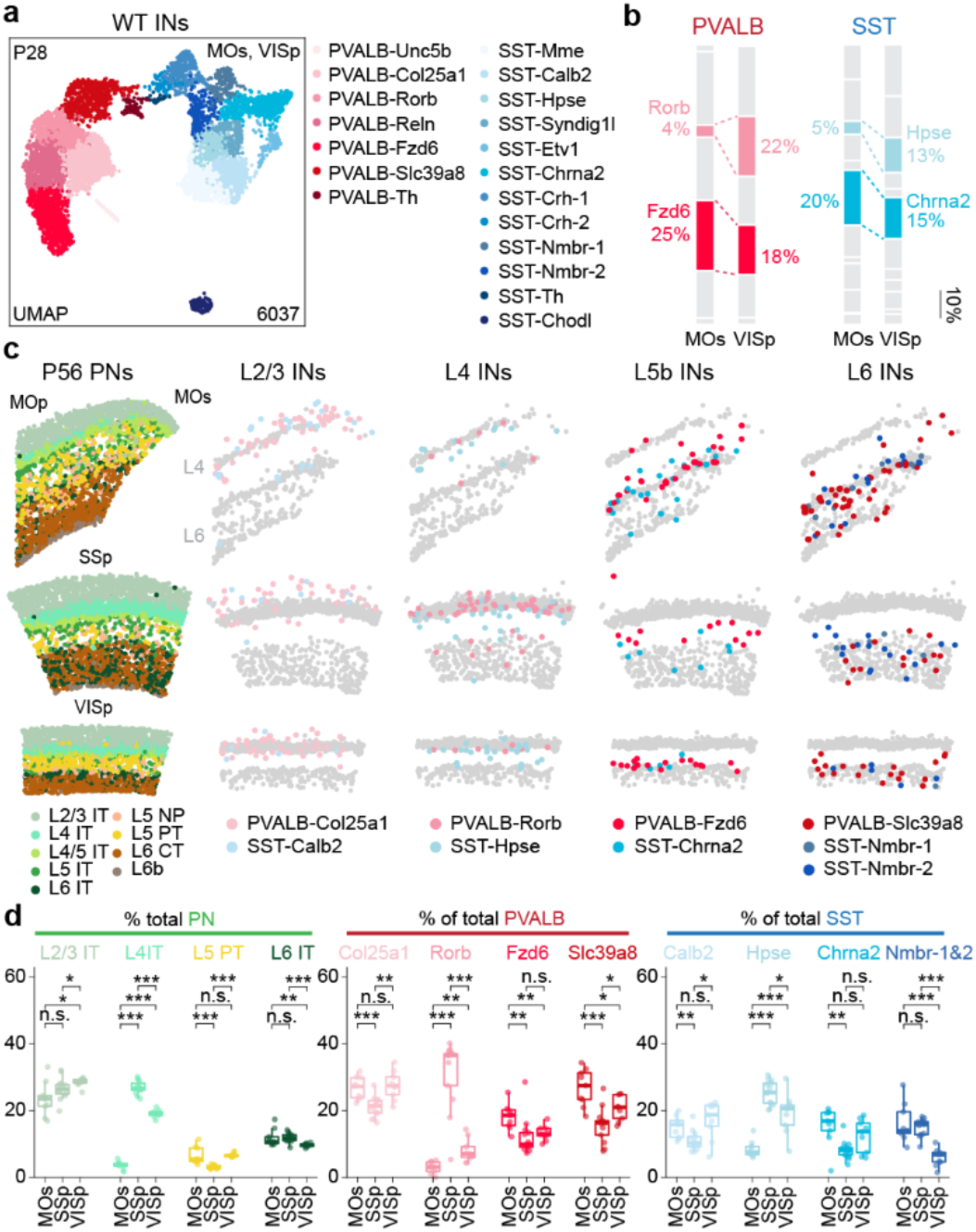
PVALB and SST interneuron subtypes are quantitatively coupled to the surrounding pyramidal neuron population. **a,** Uniform manifold approximation and projection (UMAP) visualization of snRNA-seq data from P28 cortical interneurons^11^, depicting 7 PVALB and 12 SST subtypes. Data collected from MOs (ALM) and VISp cortices. **b,** Subtype composition within PVALB or SST populations in MOs and VISp from snRNA-seq data. **c,** Representative MERFISH spatial map of coronal brain sections (P56 mouse) from published datasets^12^, illustrating the distribution of PNs and selected interneuron subtypes across three cortical regions. **d,** Proportions of selected PN, PVALB, and SST subtype by cortical region. Data in **d, e** are from n = 9 (MOs), 13 (SSp), and 9 (VISp) regions of interest (ROIs). Wilcoxon rank-sum test without multiple-comparison corrections, n.s. not significant, p≥0.05; *p<0.05; **p<0.01; ***p<0.001. Exact sample genotypes and *P* values are provided in Supplementary Tables 2 and 3.

To determine if these transcriptomic subtypes occupy distinct spatial niches, we analyzed their distribution in a whole-brain multiplexed error-robust fluorescence in situ hybridization (MERFISH) dataset^12^. Each subtype displayed a stereotyped laminar distribution that was consistent across cortical regions (Fig. 1c, Extended Data Fig. 2b, Fig. 3a-b). This positioning was highly consistent with known connectivity; for instance, L4-targeting subtypes (PVALB-Rorb, SST-Hpse) were enriched in and near L4, while L5b-targeting subtypes (PVALB-Fzd6, SST-Chrna2) were both co-localized in L5b, though they converge onto L5b PNs via distinct subcellular targeting: PVALB-Fzd6 extended their axons laterally within L5b, whereas SST-Chrna2 projected their axons to L1 and targeted the dendrites of L5b pyramidal tract (PT) neurons^4,13^ (Extended Data Fig. 1f,g). Together, these observations suggest that interneuron subtypes occupy stereotyped laminar niches that position them for highly specific interactions with distinct PN types.

## Interneuron subtype abundance changes regionally in accord with local pyramidal neuron composition across cortical regions

Although the isocortex contains a shared set of neuronal subtypes^2,3^, analysis of the MERFISH dataset (Extended Data Fig. 2i) revealed that the relative abundance of interneuron subtypes quantitatively changes according to the varying proportions of local pyramidal neuron (PN) subtypes. This relationship is most evident in L4, where intratelencephalic (IT) neurons expand from just 3% of PNs in MOs to ∼25% in primary sensory cortices. Correspondingly, the proportion of L4-targeting interneurons such as SST-Hpse and PVALB-Rorb showed a similar dramatic increase in primary sensory areas^14^ (Fig. 1c,d, Extended Data Fig. 2a). For PVALB-Rorb in particular, its relative proportion increased more than tenfold in SSp, and nearly threefold in VISp, directly contributing to the higher overall PVALB interneuron density observed in primary sensory cortices, whereas the overall density of SST interneurons remains relatively stable across cortical regions^15^ (Extended Data Fig. 2d-f). Conversely, interneuron subtypes associated with L5b PNs, SST-Chrna2 and PVALB-Fzd6, precisely mirrored the shifting proportion of L5 PT neurons across cortical regions, peaking in motor cortex (Fig. 1c,d, Extended Data Fig. 2a). These patterns were corroborated by snRNA-seq data (Fig. 1b, Extended Data Fig. 1e) and further confirmed for these SST subtypes using targeted genetic labeling (Extended Data Fig. 2g-h). Even more subtle regional differences followed this rule, such as a reduced proportion of L6 interneurons, PVALB-Slc39a8 and SST-Nmbr1/2, in VISp that correlated with the smaller L6 IT population in that region (Fig. 1d, Extended Data Fig. 1e). Full subtype-by-layer breakdowns for MOs, SSp and VISp regions are provided (Extended Data Fig. 2c, 3b,c). Thus, local interneurons appear to calibrate their laminar positioning and subtype proportions through interactions with resident PN subtypes, ultimately shaping the highly specific microcircuitry seen in the adult cortex.

## Pyramidal neuron identity controls interneuron subtype composition

The tight coupling between PN and interneuron proportions raises a fundamental question of causality: does this relationship arise from parallel, independently regulated developmental programs, or do local PNs actively instruct the composition of neighboring interneuron populations? Building on foundational studies showing that PN identity can shape interneuron laminar distribution and modulate inhibitory synaptic properties^16–22^, we tested whether PN identity affects interneuron development at subtype-specific level using the *Fezf2* knockout (KO) mouse model. *Fezf2* is an ideal target as its deletion causes a fate switch of L5 subcerebral projection neurons into callosal projection neurons, offering a clean system to study how interneurons respond to a change in their local PN partners^17,23,24^ without being confounded by direct effects on interneuron themselves, which at no time during development express *Fezf2* ^25^.

Consistent with previous reports, our analysis^26^ confirmed that deep-layer PNs in *Fezf2* KO cortex undergoes a broad shift towards an IT identity. The most drastic change is a complete loss of L5b near-projecting (NP) and PT neurons, and a corresponding increase in L6 IT neurons. While the L6 corticothalamic (CT) population was also profoundly altered^27^, we refer to these cells as L6 CT* in *Fezf2* KO to reflect their partial resemblance (Extended Data Fig. 4a-d).

To determine how interneuron populations respond to this PN fate switch, we performed snRNA-seq on *Dlx5/6-Cre*-labeled interneurons from both control and *Fezf2* KO cortices at P20. Remarkably, the changes in interneuron composition closely mirrored changes in their PN partners and reflected the known connectivity for the affected subtypes. Two interneuron subtypes associated with L5b, PVALB-Fzd6 and SST-Chrna2, were reduced by approximately 80% in *Fezf2* KO cortex. Conversely, subtypes residing in L6, PVALB-Slc39a8 and SST-Nmbr-1/2, nearly doubled in proportion (Fig. 2a,b). This shift aligns with experimentally confirmed connectivity for SST subtypes, with SST-Chrna2 preferentially innervating L5 PT, whereas SST-Nmbr-1/2 preferentially targeting deep-layer IT neurons^4^. Additionally, L6 PVALB-Th interneurons appear to be lost in the *Fezf2* mutant (Extended Data Fig. 4e,f), likely due to the loss of L6b, which they predominantly innervate (data not shown).

**Figure 2.**
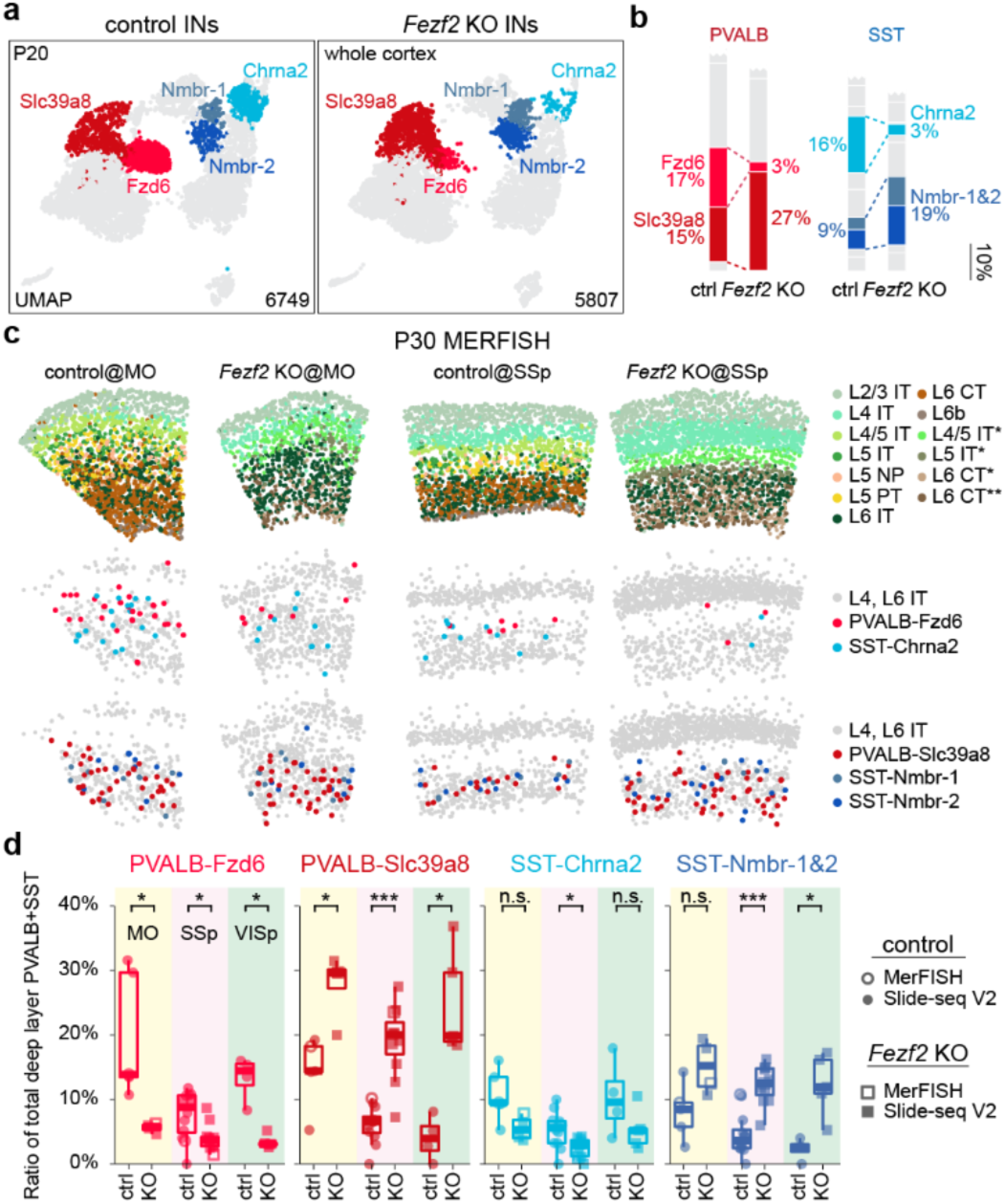
*Fezf2* KO changes the proportion of deep-layer PVALB and SST interneuron subtypes. **a,** UMAP of snRNA-seq data from P20 control and *Fezf2* KO cortical interneurons, highlighting five deep-layer interneuron subtypes with altered proportions in the mutant condition. **b,** Proportions of PVALB and SST subtypes in control and *Fezf2* KO cortices based on snRNA-seq data, with selected subtypes highlighted. **c,** Representative MERFISH spatial maps of MO and SSp coronal sections from P30 control and *Fezf2* KO mice. **d,** Proportions of selective PVALB and SST subtypes within L5/6, combining data from MERFISH and Slide-seq. Sample sizes are presented as n = mice (from X independent experiments/ROIs). MO_ctrl: n=4 (5); MO_*Fezf2* KO: n=4 (4); SSp_ctrl: n=8 (12); SSp_*Fezf2* KO: n=8 (11); VISp_ctrl: n=4 (4); VISp_*Fezf2* KO: n=2 (5). All samples from mice aged 4-6 weeks (including one published Slide-seq dataset^42^: puck 200306_02). Wilcoxon rank-sum test, n.s. not significant, p≥0.05; *p<0.05; **p<0.01; ***p<0.001. Exact sample genotypes and *P* values are provided in Supplementary Tables 2 and 3.

To confirm these quantitative changes and map the spatial distribution of deep-layer interneuron subtypes in mutant cortex, we performed MERFISH and Slide-seq spatial transcriptomic analyses on adult control and *Fezf2* KO cortices (Fig. 2c; Extended Data Fig. 5a-e). Both methods confirmed the loss of L5b and the expansion of L6 PVALB and SST interneurons in *Fezf2* mutants, with changes consistent across MOs, SSp and VISp (Fig. 2d). Furthermore, the few remaining PVALB-Fzd6 and SST-Chrna2 interneurons in the *Fezf2* KO cortices were displaced towards L6, whereas the expanded PVALB-Slc39a8 and SST-Nmbr-1/2 populations were dispersed more broadly but remained largely confined to the expanded L6 in the mutant cortex (Extended Data Fig. 5f). These shifts in the number and distribution of SST subtypes were further confirmed by genetic labeling of SST-Chrna2 interneurons and RNAscope *in situ* hybridization for SST-Nmbr-1/2 markers in control and *Fezf2* KO cortices (Extended Data Fig. 4i,j).

We next assessed how the PN fate-switch affect the molecular and cellular properties of the altered interneuron populations – specifically, how the remaining L5b interneurons persisted and adapted in the absence of their normal partners, as well as the extent to which the expanded L6 interneurons preserve their native identity. In *Fezf2* mutants, the few surviving PVALB-Fzd6 and SST-Chrna2 interneurons lost many of their subtype-defining transcriptomic features, likely reflecting improper maturation in the absence of their normal PN partners. In contrast, SST-Nmbr-1/2 interneurons acquired novel transcriptomic signatures (Extended Data Fig. 4g-h), accompanied by marked morphological remodeling, including more elongated dendrites along the cortical column, and fewer axonal projections to L1, relative to controls (Extended Data Fig. 4k). The expanded L6 PVALB-Slc39a8 interneurons, however, showed no significant transcriptomic changes. Consistent with the general stability of PVALB interneuron transcriptomes across cortical regions, these results further suggest that PVALB interneurons are less sensitive to subtle transcriptomic changes in their associated PN subtypes, except when PN identity is entirely abrogated.

Finally, beyond the changes in the deep-layer interneurons described above, previous studies have reported an upward laminar shift of PVALB and SST interneurons in the *Fezf2* KO cortex^17^. Consistent with this, our MERFISH analyses reveals that this phenomenon is due to a spatial redistribution—rather than a change in number—of specific mid-layer subtypes, such as PVALB-Rorb (and, to a less extend, PVALB-Reln) and SST-Hpse (Extended Data Fig. 6).

Thus, by manipulating pyramidal neuron identity using the *Fezf2* mutant, we demonstrate that the subtype composition and laminar distribution of PVALB and SST interneurons are non-autonomously affected by their specific PN partners.

## *Bax* deletion rescues the SST, but not PVALB, interneuron phenotype in *Fezf2* mutants

A developmental wave of apoptosis, modulated by PNs, normally refines cortical interneuron numbers around P7-P9^28–30^. To determine if the shift in interneuron subtype proportions in *Fezf2* mutants is driven by an exaggeration of this programmed cell death versus a failure in fate specification, we blocked apoptosis by conditionally removing the pro-apoptotic gene *Bax* from PVALB and SST interneuron precursors well before the onset to this apoptosis period.

To first establish the baseline effects of inhibiting apoptosis, we removed *Bax* in a wild-type background using the *Nkx2.1-Cre* driver line. RNAscope *in situ* hybridization revealed that this manipulation increased the number of PVALB (∼22%) and SST (∼31%) interneurons without significantly altering their overall laminar distribution (Extended Data Fig. 7b,c,f,h). Consistent with this, quantification of the *Nkx2.1-Cre*-labeled interneurons showed a ∼36% increase in the total population (Extended Data Fig. 7d-e). The apparent larger effect size with this quantification likely reflects the known sparing of superficial interneurons not targeted by the transgenic *Nkx2.1-Cre* driver^31^ (Extended Data Fig. 7a) that still undergoes apoptosis. Applying the same *Bax* deletion on the *Fezf2* KO background increased the number of PVALB and SST interneurons by a similar magnitude (Extended Data Fig. 7c,g), consistent with previous reports that *Fezf2* mutant does not increase overall interneuron cell death^17^.

With the baseline effects established, we next used snRNA-seq to directly test how preventing apoptosis affects interneuron subtypes composition in *Fezf2* mutants. To this end, we profiled PVALB and SST interneurons across four different genotypes: control, *Bax* cKO, *Fezf2* KO, and *Fezf2* KO_*Bax* cKO at P14, an age when apoptosis in interneurons has normally concluded. After computationally removing a small population of non-cortical-like interneurons unique to the *Bax* cKO samples (Extended Data Fig. 7i-j), the remaining datasets integrated well across all conditions (Fig. 3a), with biological replicates for the two *Bax* cKO genotypes showing high consistency (Extended Data Fig. 7k). In line with no overall laminar distribution in the wild-type background noted above, , *Bax* deletion did not substantially alter the proportions of most subtypes. Strikingly, however, *Bax* deletion in the *Fezf2* KO background had a differential effect: it robustly rescued the loss of SST-Chrna2 interneurons, restoring their proportion close to control levels, but failed to rescue the depleted PVALB-Fzd6 population (Fig. 3b).

**Figure 3.**
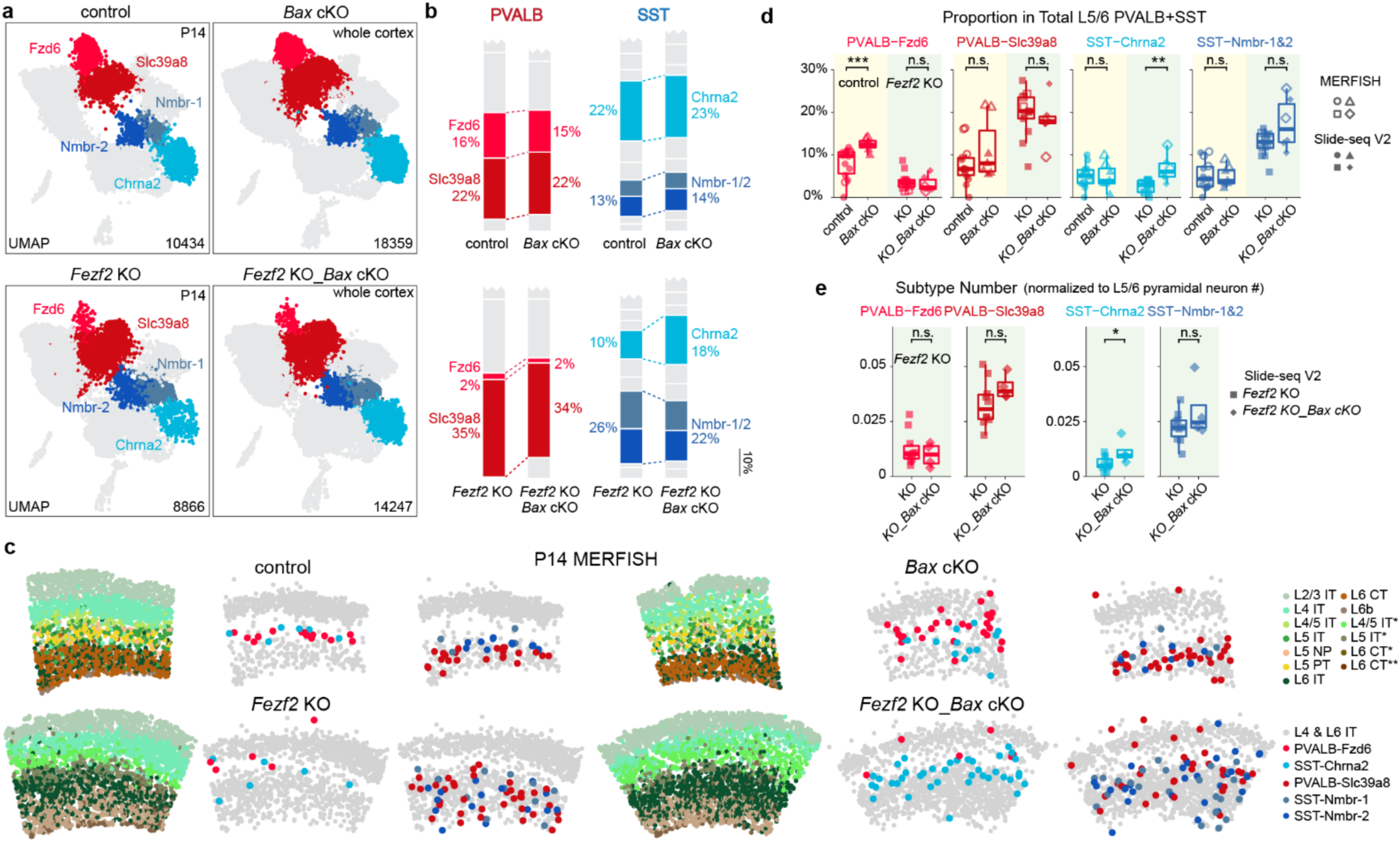
Preventing cell death rescues SST but not PVALB interneuron subtypes in *Fezf2* mutants. **a,** UMAP visualization of snRNA-seq of sorted interneurons from control, *Bax* cKO, *Fezf2* KO and *Fezf2* KO_*Bax* cKO mice at P14. Control, n = 1 (1); *Bax* cKO, n = 2 (2); *Fezf2* KO, n = 1 (1); *Fezf2* KO; *Bax* cKO, n = 4 (2). **b,** Proportion of deep-layer PVALB and SST interneuron subtypes in snRNA-seq dataset. **c,** MERFISH spatial map of coronal sections from SSp cortices of P14 mice for each genotype. **d,** Proportion of selected PVALB and SST subtypes within L5/6 of SSp, combining data from MERFISH and Slide-seq. Control: n=10 (14); *Bax* cKO: n=4 (7); *Fezf2* KO: n=10 (13); *Fezf2* KO_*Bax* cKO: n=4 (6). **e,** Number of selected PVALB and SST subtypes normalized to the number of deep-layer PNs based on Slide-seq data. *Fezf2* KO: n=7 (10); *Fezf2* KO_*Bax* cKO: n=3 (4). For **d-e**, Age range: P14-37, with one *Fezf2* KO mouse at 6-weeks old. Sample sizes are presented as n = mice (from X independent experiments/ROIs). Wilcoxon rank-sum test, n.s. not significant, p≥0.05; *p<0.05; **p<0.01; ***p<0.001. Exact sample genotypes and *P* values are provided in Supplementary Tables 2 and 3.

To confirm our snRNA-seq findings with spatial transcriptomics, we performed MERFISH and Slide-seq analyses on SSp cortices across the four genotypes (Fig. 3c, Extended Data Fig. 8a,c). To maximize statistical power, we combined data from relevant ages (P14 to adult) since interneuron composition is stable after P14^15^ and pooled control genotypes (wild-type and heterozygous) (Extended Data Fig. 8d), focusing our analysis only on deep-layer interneurons that can be robustly targeted by the *Nkx2.1-Cre* driver. These analyses corroborated our sequencing results. *Bax* cKO in control conditions does not substantially alter the proportion of interneuron subtypes, apart from a slight increase in PVALB-Fzd6 (Fig. 3d, Extended Data Fig. 8e). In contrast, both spatial transcriptomic approaches confirmed that the compound loss of *Bax* results in the rescue of SST-Chrna2 population, doubling its number relative to deep-layer PNs to match control levels, but failed to prevent the loss of PVALB-Fzd6 in *Fezf2* mutant (Fig. 3d,f, Extended Data Fig. 8b).

Together, these results reveal PNs provide an essential pro-survival signal for SST-Chrna2 interneurons, whose loss in *Fezf2* mutant is rescued by blocking apoptosis. In contrast, the loss of PVALB-Fzd6 must occur through a distinct, *Bax*-independent mechanism, suggesting that re-specification of their partner PNs alters their fate specification.

## Pyramidal neurons instruct the fate of PVALB interneuron subtypes

To seek direct evidence for a fate switch in PVALB-Fzd6 interneurons, we performed snRNA-seq at P7 of control and *Fezf2* KO condition to capture the cellular states just before their terminal differentiation. By this stage, PVALB-Fzd6, as well as SST-Chrna2 populations, have already begun to decline in *Fezf2* KO but remained sufficiently abundant for robust analysis (Extended Data Fig. 9a-b).

RNA velocity analysis^32^, which infers transcriptional momentum of individual cells based on the ratio of spliced and unspliced mRNAs, showed a pronounced flow of PVALB-Fzd6 interneurons towards a PVALB-Slc39a8 identity in the mutant, rather than controls where these cell types diverge during this period (Fig. 4a). To quantify this predicted fate switch longitudinally, we applied optimal transport mapping^33^ across P7 and P14 datasets. This analysis generated a fate assignment for each P7 subtype, predicting that ∼80% of PVALB-Fzd6 interneurons acquire a PVALB-Slc39a8 identity by P14 in the *Fezf2* mutant. Moreover, analysis of their origin from P14 back to P7 further corroborated this, showing that the expanded PVALB-Slc39a8 population in the mutant at P14 was a mix, originating from both normal P7 precursors (∼50%), as well as from P7 PVALB-Fzd6 cells (∼20%). In control conditions, by contrast, subtypes faithfully mapped between corresponding counterparts across age in both directions (Fig 4b). Thus, both analyses revealed a substantial, PN-dependent fate switch with regards to the PVALB lineage.

**Figure 4.**
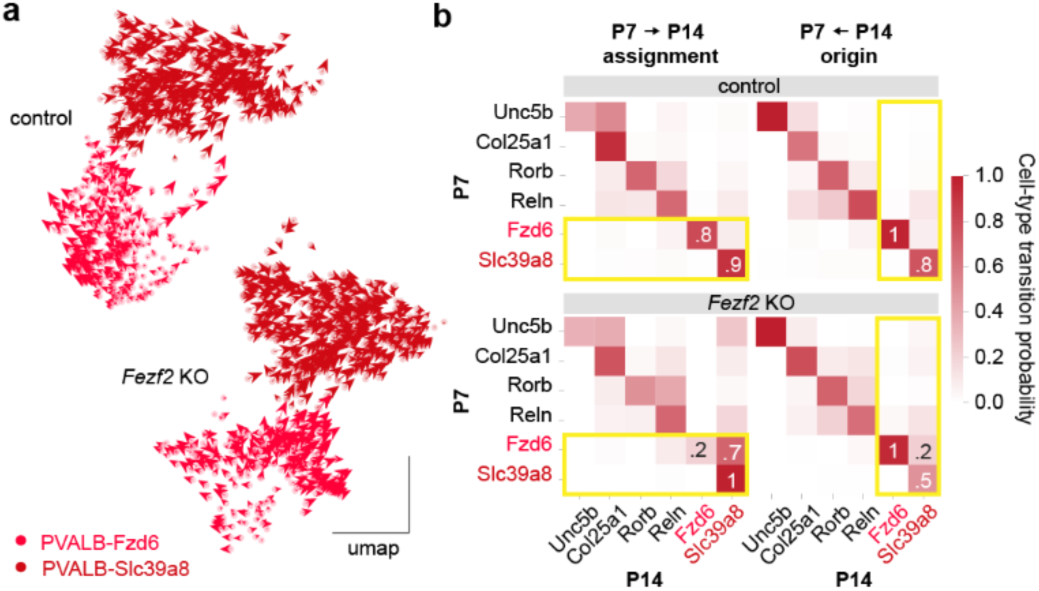
*Fezf2* deletion drives a fate switch in PVALB interneurons. **a,** RNA velocity analysis of snRNA-seq data on cortical interneurons collected from P7 control and *Fezf2* KO cortices showing transcriptomic dynamics of PVALB-Fzd6 and PVALB-Slc39a8 interneurons. **b,** Optimal transport analysis mapping lineage relationships between P7 and P14 subtypes in control and *Fezf2* KO conditions. The descendant map (left) shows the predicted assignment of P7 subtypes to P14 fates; the ancestor map (right) traces the developmental origin of P14 subtypes back to P7.

Interestingly, the same analyses revealed an indication of plasticity in the SST lineage, albeit to a much less extent. RNA velocity indicated a subtle shift of SST-Chrna2 transcriptomic dynamics of SST-Chrna2 towards SST-Nmbr-1 in mutant condition. Consistently, optimal transport predicted that ∼10% of P7 SST-Chrna2 interneurons transform into SST-Nmbr-1 by P14, accounting for ∼7% of the expanded SST-Nmbr-1 population in the *Fezf2* KO (Extended Data Fig. 10c,f).

Gene set analysis and cross-age comparisons further corroborated these distinct outcomes for PVALB and SST interneurons. Using control data as a reference, we defined marker genes (ID genes) that are differentially expressed between L5b and L6 subtypes within PVALB and SST classes. In *Fezf2* mutants, P7 PVALB-Fzd6 interneurons downregulated their expected ID genes, while upregulating those of PVALB-Slc39a8 subtype. Moreover, they were arrested in a more immature state, possessing a transcriptomic profile similar to PVALB-Fzd6 interneurons at P2 (Extended Fig. 10a,b). In contrast, SST-Chrna2 interneurons in *Fezf2* mutant largely retained their core ID genes. Although they also appeared less mature, their overall transcriptomic state remained closer to control SST-Chrna2 interneurons at P7 (Extended Data Fig. 10d,e).

Finally, despite the respecification of PN identity already apparent in *Fezf2* mutants at P2 (Extended Data Fig. 9c-e), PVALB and SST interneurons showed no overt changes in their number or identity at this age, based on both snRNA-seq and MERFISH data (Extended Data Fig. 9f-i). This indicates that the shift in subtype composition arise only after interneurons enter the cortex and interact with PNs.

Taken together, these findings indicate that PNs sculpt their associated PVALB and SST interneuron populations through two distinct mechanisms: directed differentiation or pro-survival (Fig. 6). While PVALB interneurons exhibited high plasticity, with PNs instructively guiding their fate, SST subtypes are more restricted in their fate, with PNs primarily promote the survival of SST subtypes.

## Pyramidal neurons orchestrate interneuron development through a combination of contact-mediated and secreted signals

To investigate the nature of the signaling molecules from L5 PT/NP neurons that direct interneuron development, we performed two complementary perturbation experiments in *Fezf2*-expressing neurons. Neuronal activity was suppressed postnatally through overexpression of Kir2.1, or VAMP2-dependent vesicular release was disrupted via tetanus toxin (TeNT) expression upon the onset of *Fezf2* expression.

Silencing activity did not alter the relative proportions of PVALB and SST interneuron subtypes. In contrast, blocking vesicular release partially recapitulated the *Fezf2* knockout phenotype, leading to a reduction in L5b-enriched PVALB-Fzd6 alongside an increase in L6-enriched PVALB-Slc39a8 interneurons. L5b and L6 SST interneurons showed a similar trend in their relative abundance, but the effects were much more modest (Fig. 5a,b, Extended Data Fig. 11a-e). Despite a milder effect on cell number, the remaining PVALB-Fzd6 cells in TeNT-treated brains showed a more pronounced transcriptomic shift than those in the *Fezf2* knockout, suggesting that they are likely stalled in an unstable, intermediate state, potentially because the surrounding pyramidal neurons are silenced but not respecified into the L6-like fate seen in the full knockout (Extended Data Fig. 11f).

**Figure 5.**
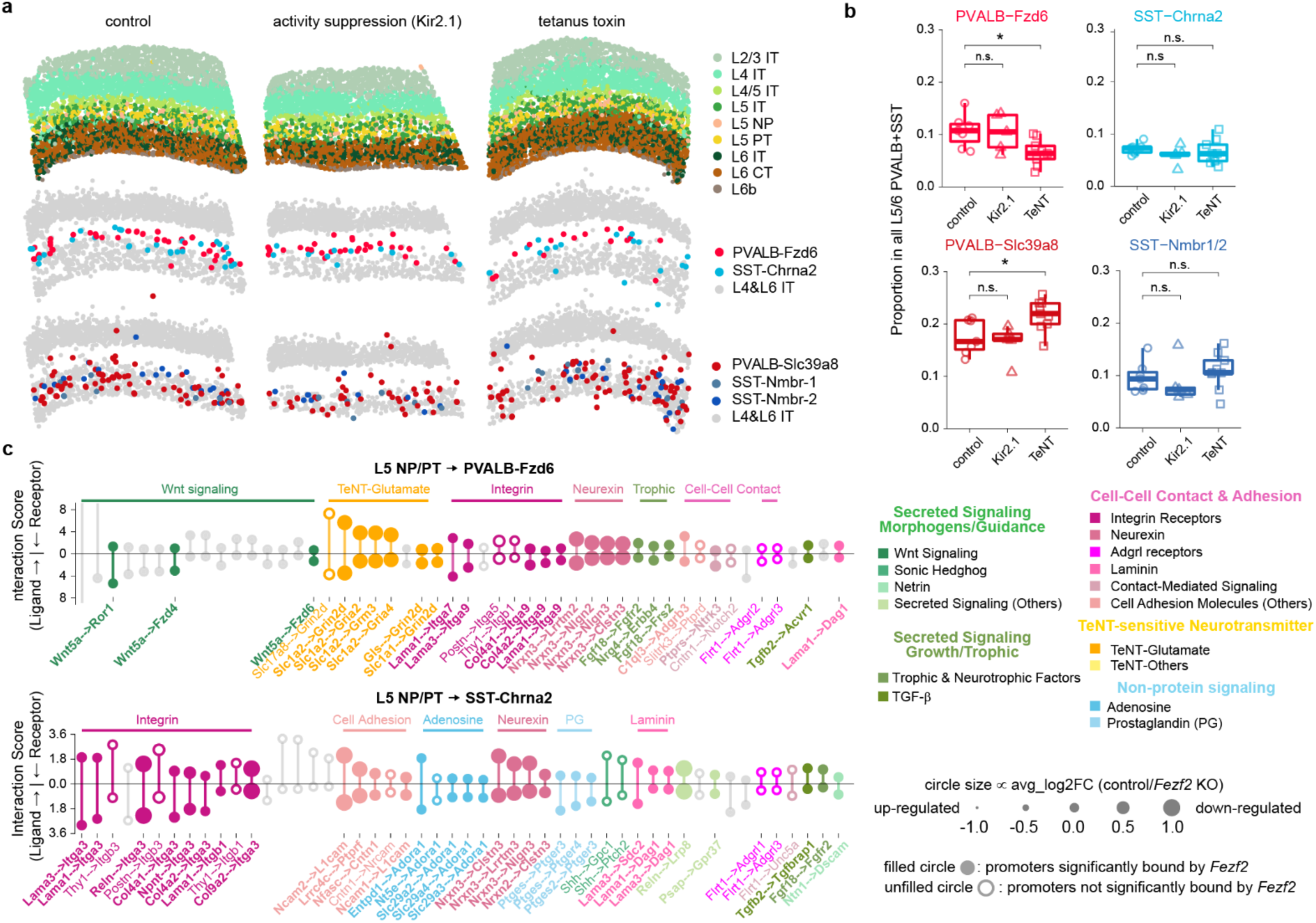
PN-to-interneuron signaling is largely voltage-gate activity-independent and mediated by distinct molecular pathways in PVALB and SST subtypes. **a,** Representative MERFISH spatial maps of SSp region showing the number and spatial location of selected PVALB and SST interneuron subtypes in control, activity silencing (Kir2.1) or blocking vesicular release (TeNT) conditions. **b,** Boxplots showing the proportions of selected deep-layer PVALB and SST subtypes across conditions based on MERFISH data. Sample sizes are presented as n = mice (from X ROIs). control: n=5 (7); Kir2.1: n=3 (5); TeNT: n=5 (9). Age range: P14-22. Wilcoxon rank-sum test, n.s. not significant, p≥0.05; *p<0.05. Exact sample genotypes and *P* values are provided in Supplementary Tables 2 and 3. **c,** Lollipop diagrams of top 50 candidate ligand-receptor pairs predicted to mediate PN-interneuron interactions between L5b PNs and L5b PVALB versus SST interneurons. Line length represents the interaction score, with contributions partitioned across the midline according to relative ligand expression in L5b PNs versus receptor expression in L5b interneurons. Circle size indicates downregulation (large) or upregulation (small) of the ligand in *Fezf2* KO. Filled circle signifies direct binding of the *Fezf2* transcription factor to the ligand’s gene, based on ChIP-seq data. See **Supplemental Tables 4 and 5** for a complete list of the top 100 ligand-receptor pairs.

These experiments allow us to parse the underlying signaling mechanisms. The lack of a Kir2.1 phenotype argues against a major role for activity-dependent processes, especially dense-core vesicle release, which usually requires high-frequency firing. The partial TeNT phenotype, however, points to a role for spontaneous VAMP2-dependent release. Crucially, because this effect is much milder than the full *Fezf2* knockout, our findings indicate that VAMP2-independent signals must be another major drive in sculpting the local interneuron population.

Guided by these findings, we performed a targeted bioinformatic to identify candidate ligand-receptor pairs that mediate this interaction. To move beyond generic prediction algorithms, we applied two strict criteria: (1) the ligand or receptor must show subtype-specific expression in the presynaptic PN and postsynaptic interneuron, respectively and (2) the PN-derived ligand must be downregulated in the *Fezf2* KO. This analysis revealed distinct candidate pathways for each interneuron class. For SST-Chrna2, top candidates were primarily cell-adhesion molecules, particularly those involving integrin receptors, suggesting their survival is supported by a combinatorial code of cell-cell contacts (Fig. 5c, Extended Data Fig. 12a). For PVALB-Fzd6, however, the analysis pointed overwhelmingly to non-canonical Wnt signaling. The ligand Wnt5a met all criteria: it is specifically expressed in L5b PNs at the relevant age (∼P4)^34^, its expression is dependent on *Fezf2*, and ChIP-seq data^35^ indicates it is a direct transcriptional target of *Fezf2* (Fig. 5c, Extended Data Fig. 12a, b). Thus, our findings indicate that pyramidal neurons utilize a diverse repertoire of signaling mechanisms, including both tetanus toxin-sensitive and -insensitive pathways, to ensure the precise assembly of local circuits (Fig. 6).

**Figure 6.**
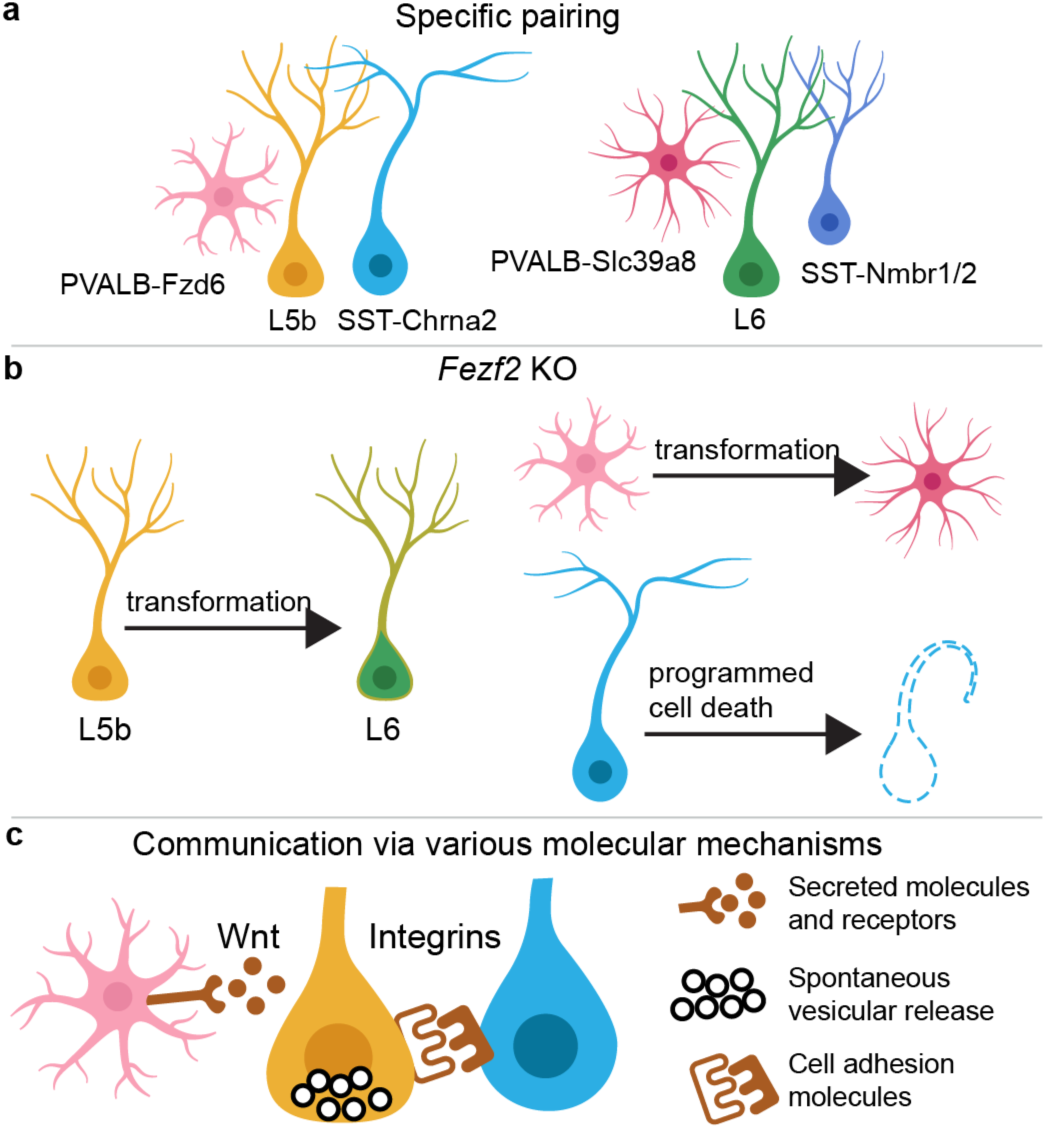
A model for the regulation of interneuron subtype composition by local pyramidal neurons. In the developing cortex, the subtype composition of local interneurons is quantitatively coupled to that of the surrounding pyramidal neuron (PN) population. In a *Fezf2* mutant where deep-layer PNs are altered, this balance is maintained through two divergent mechanisms: SST interneurons adjust their numbers via programmed cell death, while PVALB interneurons undergo a cell fate transformation. This regulation is mediated by distinct PN-derived signals; the PVALB fate switch depends on spontaneous, tetanus toxin (TeNT)-sensitive vesicular release, while SST interneuron survival relies on TeNT-insensitive signals. Our analysis identifies the Wnt5a pathway as a top candidate for regulating PVALB interneurons and integrin-mediated signaling for SST interneurons.

## Discussion

Our findings reveal that pyramidal neurons (PNs) are not passive scaffolds but active sculptors of cortical inhibitory circuits, employing distinct mechanisms matched to the intrinsic plasticity of their interneuron partners. We show that PNs instructively guide the fate PVALB subtypes while primarily promoting the survival of SST subtypes which, unable to adapt by changing fate, are eliminated by apoptosis.Consistent with this, preliminary snMultiome data revealed potential difference in their epigenetic landscapes during development. We found that PVALB interneurons at P7 exhibit a “poised” chromatin state, where genes defining an alternative fate are just as accessible as their own. In contrast, SST interneurons appear more epigenetically committed, with their own identity genes showing significantly higher accessibility (data not shown). A compelling explanation for this divergence is the developmental timing of the subtypes involved. PVALB interneurons, which mature later^36–38^, appear to retain a more malleable identity that allows for PN-guided fate changes. In contrast, SST subtypes are committed to their fate earlier, making them dependent on PN-derived signals for survival rather than re-specification. However, this distinction is not absolute, the potential of a small fraction of SST interneurons to change fate suggests plasticity is a continuum, likely reflecting a dynamic interplay between cell-intrinsic programs and the extrinsic, PN-derived cues that ultimately shape the final circuit.

While our data demonstrates a uniquely powerful and subtype-specific influence exerted by PNs, they act within a complex developmental environment that includes other factors such as glia and the extracellular matrix^39–41^. Our work, however, establishes PN-interneuron interaction as a central player in sculpting the fine-grained composition of the local inhibitory network. This provides a new framework for understanding how the immense diversity of cortical interneurons is generated and precisely matched to the principal neurons they serve, ensuring the formation of a balanced and functional cortical circuit.

## Supporting information

Supplementary file list

---

**Extended Data Fig. 1.**
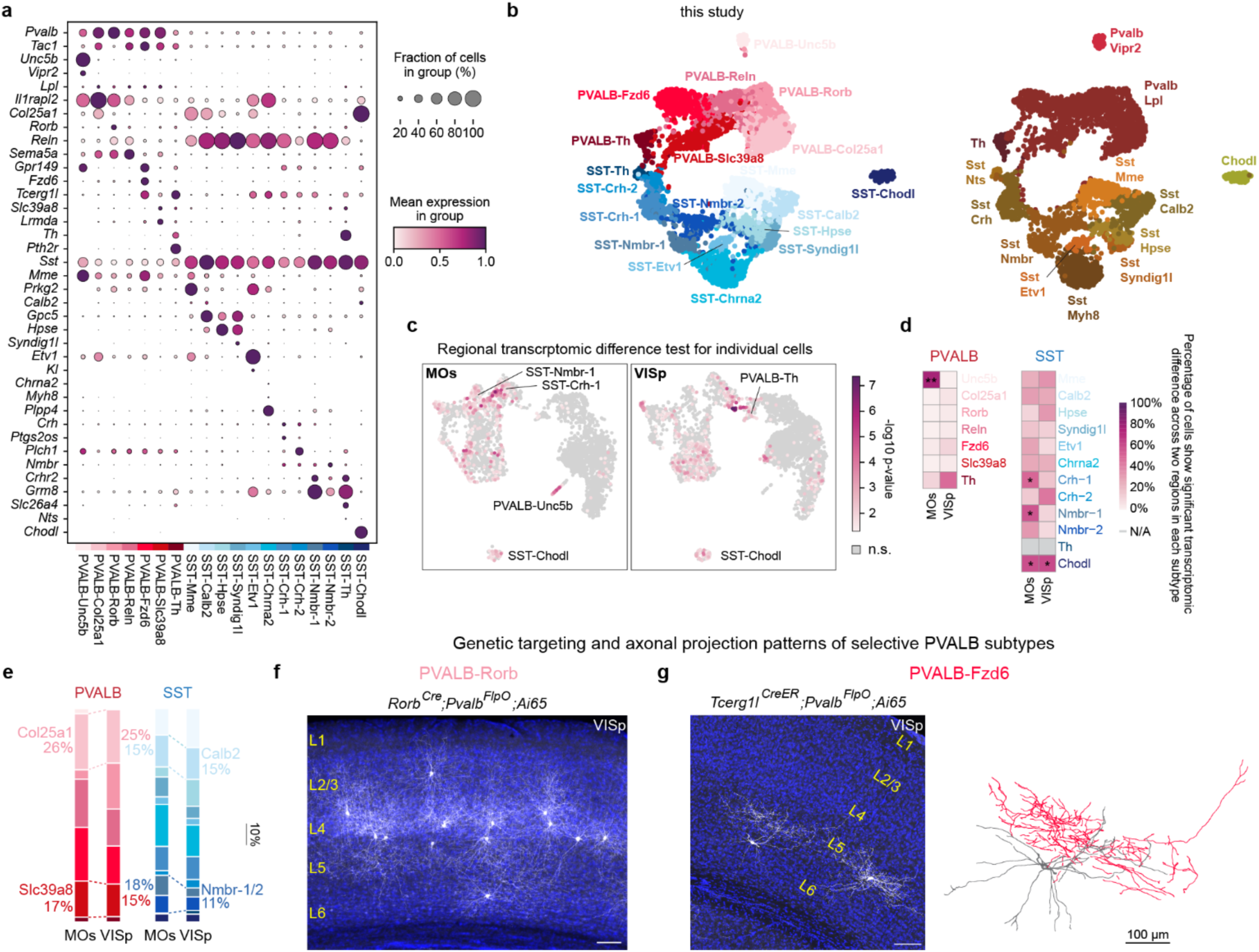
Characterization and regional transcriptomic analysis of PVALB and SST interneuron subtypes. **a,** Dot plot showing expression of marker genes for each PVALB and SST interneuron subtype. **b,** Correspondence between subtypes identified in this study and previously defined supertypes^3^. **c,** Regional transcriptomic differences of PVALB and SST interneurons between MOs and VISp assessed with Emergene (see Methods), which calculates a *P*-value for region-specific gene signatures in individual nuclei using permutation tests. n.s., not significant. **d,** Heatmap showing the percentage of cells per subtype with significant regional differences. Subtypes with ≥50% significant cells are annotated with the median *P*-value as calculated by Emergene (*p<0.05, **p<0.01). N/A, not analyzed (cluster size <10). Exact *P*-values are provided in Supplementary Table 3. **e,** Proportions of all PVALB and SST subtypes in MOs and VISp, complete version of Figure 1b. **f,** Intersectional genetic strategy preferentially targeting PVALB-Rorb interneurons, showing their axons concentrated in L4. **g,** Intersectional genetic strategy preferentially targeting PVALB-Fzd6 interneurons, showing that these interneurons reside in L5b and extend their axons laterally within L5b. Sparse labeling achieved by low-dose tamoxifen administration allowed reconstruction of individual PVALB-Fzd6 morphologies, with one example shown (right). For **f-g**, note that both strategies have off-target labeling of L5 PT neurons specifically in SSp regions. Scale bars, 100 µm.

**Extended Data Fig. 2.**
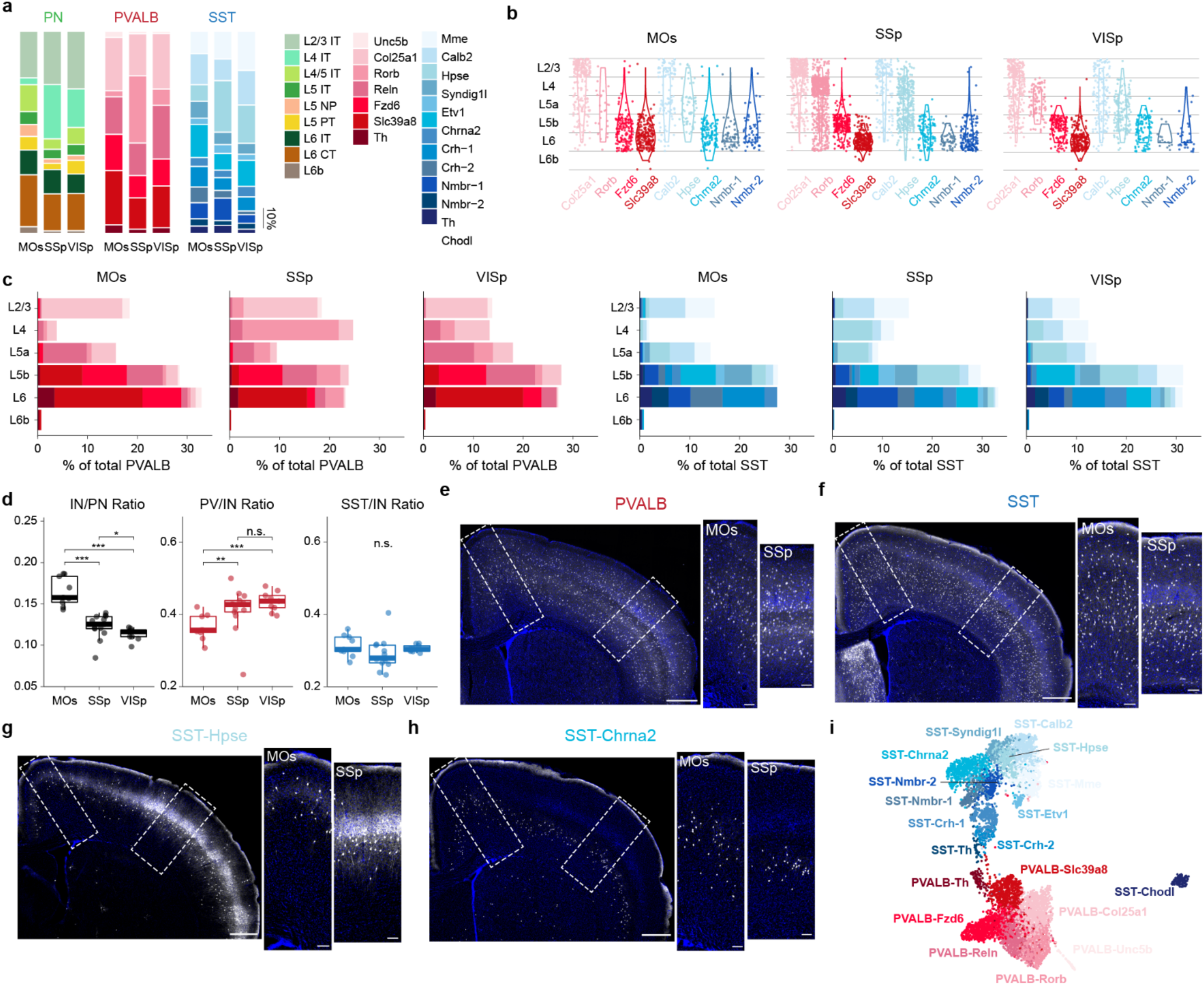
Regional differences in the proportion of different PVALB and SST interneuron subtypes. **a,** Proportion of all PN, PVALB, and SST interneuron subtypes in MOs, SSp, and VISp cortices based on MERFISH data. **b,** Violin plots showing the laminar distribution of selected PVALB and SST subtypes across three cortical regions. **c,** Bar plots showing the layer-by-layer composition of PVALB and SST subtypes in each region. **d,** Boxplots showing the ratio of interneurons (INs) to PNs and the contribution of PVALB and SST subtypes to the total interneuron pool across regions. Wilcoxon rank-sum test, n.s. not significant, p≥0.05; *p<0.05; **p< 0.01; ***p<0.001. Exact *P* values are provided in Supplementary Tables 3. **e–h**, Validation of regional interneuron densities by immunolabeling and genetic targeting. **e,** Immunolabeling of PVALB interneurons, **f,** Genetic labeling of all SST interneurons; **g,** Labeling of L4-targeting SST-Hpse interneurons with some off-target SST-Calb2 interneurons; **h,** Labeling of L5b SST-Chrna2 interneurons. Scale bars, 500 µm (main) and 100 µm (higher magnification of outlined regions on the right). Exact sample genotypes are provided in Supplementary Tables 2. **i,** UMAP of P28 snRNA-seq data re-clustered using only the genes included in the MERFISH probe set, demonstrating that this gene set is sufficient to resolve the defined subtypes.

**Extended Data Fig. 3.**
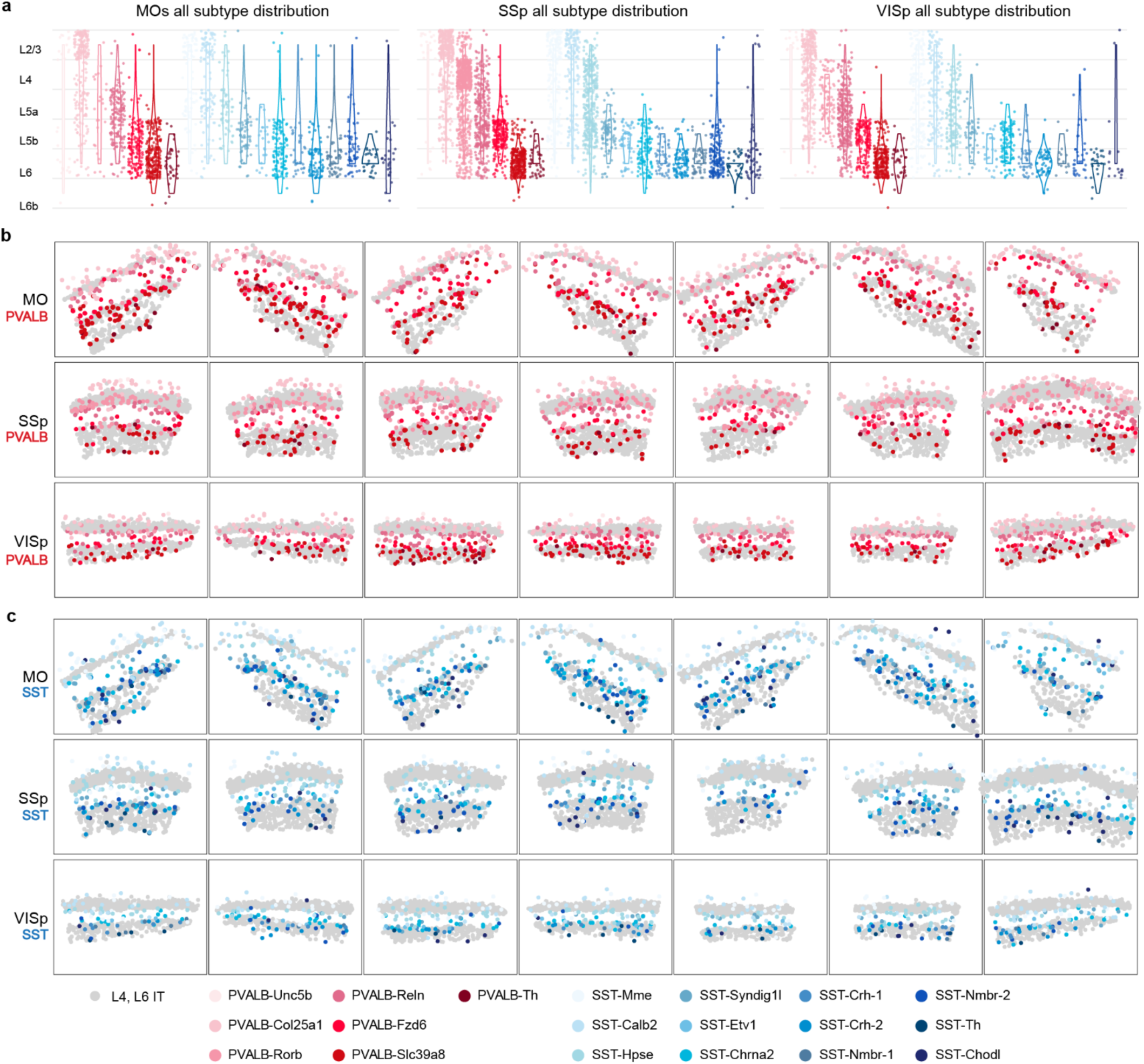
PVALB and SST interneuron subtypes show stereotyped laminar distributions. **a,** Violin plots showing the laminar distribution of all PVALB and SST subtypes across MOs, SSp, and VISp. **b-c,** Gallery of MERFISH spatial maps from additional coronal sections showing the consistent distribution of all identified PVALB and SST interneurons.

**Extended Data Fig. 4.**
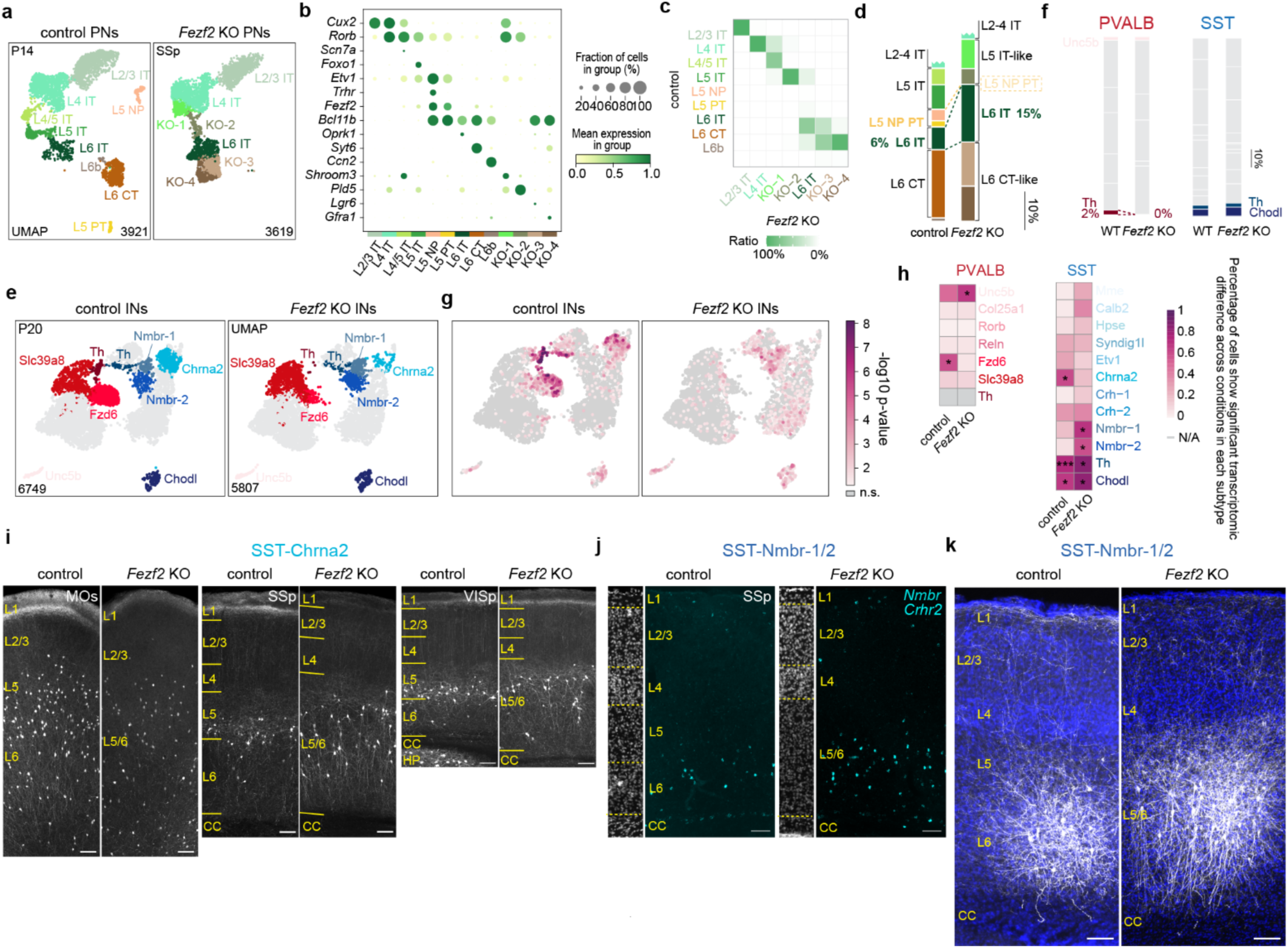
Characterization of pyramidal neuron and interneuron changes in *Fezf2* mutants. **a–d,** Analysis of PN subtypes in the P14 *Fezf2* KO SSp cortex^26^. **b,** Dot plot showing marker gene expression. **c,** Heatmap showing transcriptomic correspondence between control and KO subtypes. **d,** Quantification of deep-layer PN subtype proportions. **e–h,** Analysis of interneuron subtypes in the *Fezf2* KO. **e,** UMAP (as in Fig. 2a) highlighting all subtypes with altered proportions or transcriptomes. **f,** Bar plot (as in Fig. 2b), highlighting additional subtypes that showed changes in *Fezf2* mutants. **g,h,** Transcriptomic differences between PVALB and SST interneurons in control and *Fezf2* KO conditions (Emergene; n.s., not significant, , p≥0.05; *p<0.05; **p< 0.01; ***p<0.001). N/A: not analyzed (cluster size <10). Exact *P* values are provided in Supplementary Tables 3. **i–k**, Validation of SST subtype changes. **i,** Genetic labeling of SST-Chrna2 interneurons in different cortical regions of control and *Fezf2* mutant mice. **j,** RNAscope *in situ* hybridization against *Nmbr* and *Crhr2* mRNA transcripts, marker genes for SST-Nmbr-1 and SST-Nmbr-2 subtypes, in control and *Fezf2* KO mice at P28. **k,** Viral genetic labeled SST-Nmbr-1/2 interneurons in control and *Fezf2* mutants SSp cortices. For **i-k**, Scale bar: 100 µm. CC, corpus callosum; HP, hippocampus. Exact sample genotypes are provided in Supplementary Tables 2.

**Extended Data Fig. 5.**
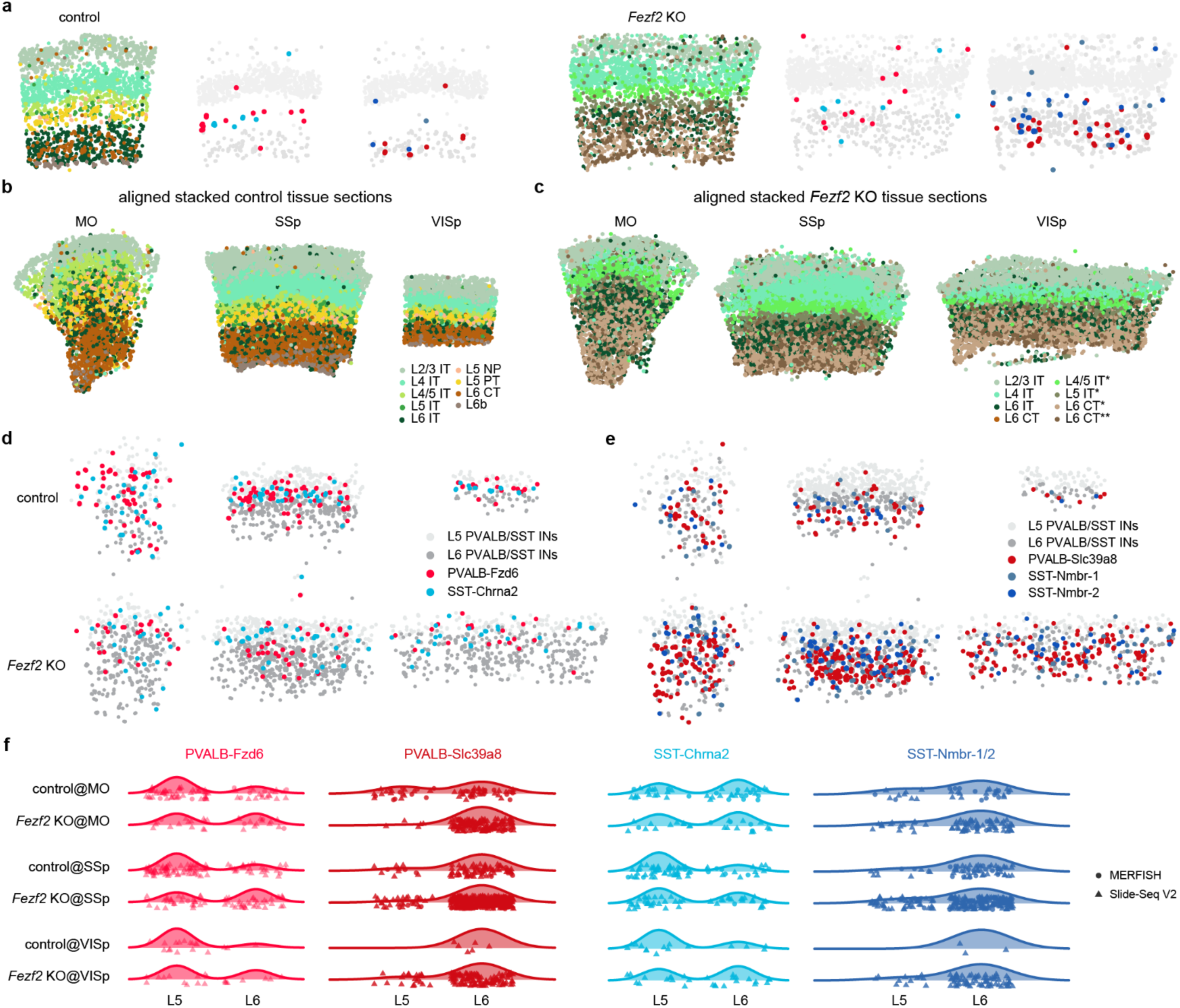
Spatial Transcriptomic analysis of cortical interneuron changes in *Fezf2* mutants. **a,** Representative Slide-seq data on a single coronal brain sections of the SSp cortex from WT (left) and *Fezf2* KO (right) mouse at P37. **b-c,** Aggregate Slide-seq data from multiple brain sections, showing the overall distribution of PNs in control (**b**) and *Fezf2* KO (**c**) cortices. Sample sizes are presented as n = mice (from X ROIs). WT control: MO, n=3 (4); SSp, n=5 (8); VISp: n=4 (4). Age: P28-37. *Fezf2* KO: MO, n=3 (3); SSp, n=5 (6); VISp, n=2 (4). Age: 4-6 weeks. **d-e,** Identification of deep-layer PVALB and SST interneuron subtypes within the aggregate control (**d**) and *Fezf2* KO (**e**) datasets. (same dataset as in **b-c**). **f,** Ridge plots showing the distribution of selected PVALB and SST subtypes in the deep layers of control and *Fezf2* KO brains, based on MERFISH data as in Fig. 2c.

**Extended Data Fig. 6.**
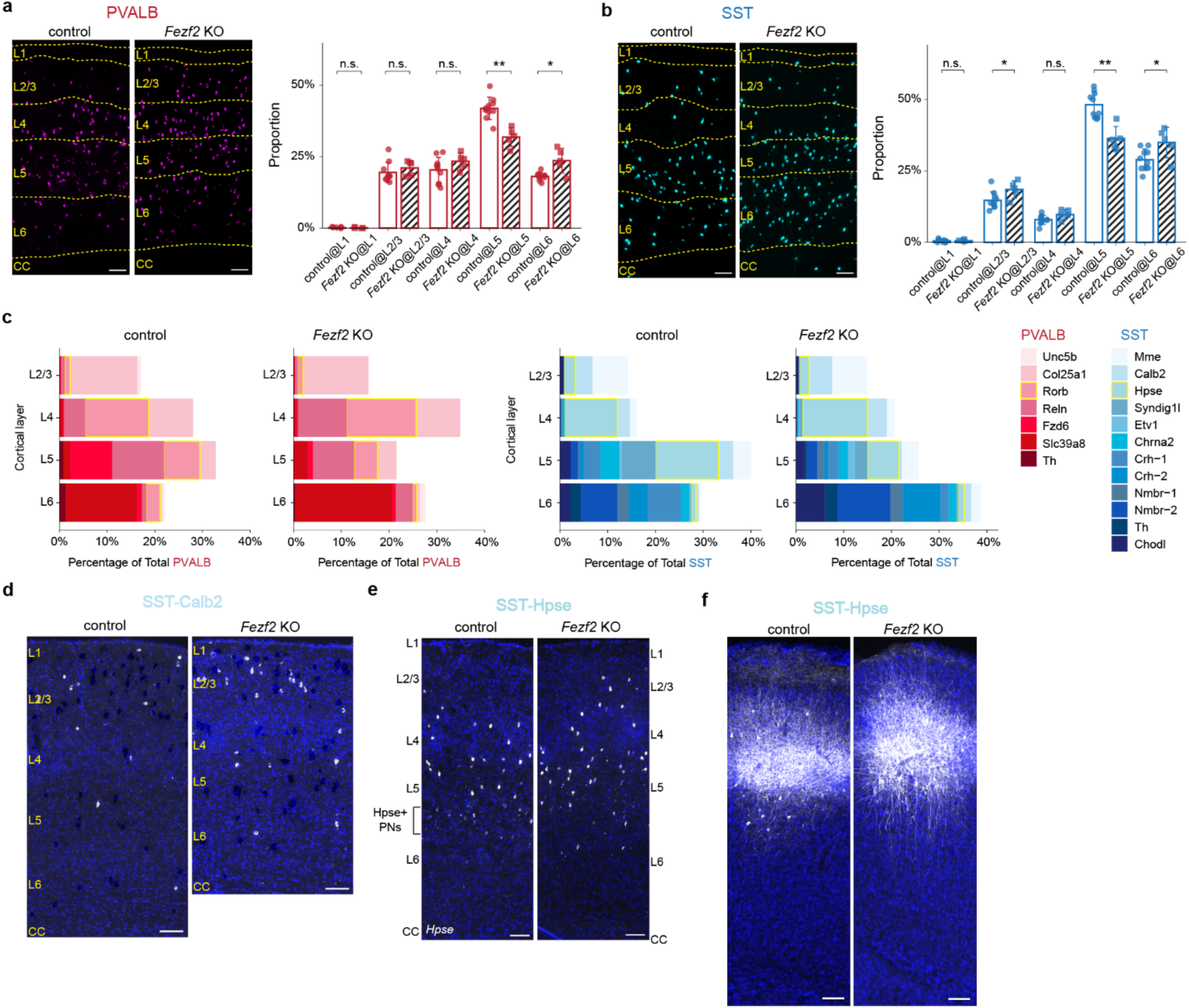
*Fezf2* deletion causes a spatial redistribution of mid-layer interneuron subtypes. **a, b,** RNAscope in situ hybridization confirms an upward laminar shift of the total PVALB (**a**) and SST (**b**) interneuron populations in the SSp of P28 *Fezf2* KO mice. Scale bar, 100 µm. Bar plots (right) quantify the proportion of cells per layer. Sample sizes are presented as n = mice (from X ROIs). control: n=5 (10), age P26-28, n=3932 PVALB, n=2280 SST. *Fezf2* KO: n=2 (5), age P27-28 n=2377 PVALB, n=1677 SST. Wilcoxon rank-sum test, n.s. not significant, p≥0.05; *p<0.05; **p< 0.01. **c,** Bar plots showing the composition of each PVALB and SST interneuron subtypes in different cortical layers of control and *Fezf2* KO brains in the SSp region, based on MERFISH dataset. control: n=2 (3), *Fezf2* KO: n=3 (3), age: P14-30. **d,** Representative RNAscope images of SST-Calb2 interneurons showing no laminar shift in *Fezf2* KO cortex (SSp). Signal shown as Calb2–Vip subtraction; note that this strategy also labels L6 SST-Chodl interneurons. **e,** RNAscope images of *Hpse* mRNA in SSp from P31 control and *Fezf2* KO brains, showing a superficial shift of SST-Hpse interneurons. Note: L5 PNs express low levels of *Hpse*. **f,** Representative images of viral genetically labeled SST-Hpse interneurons in control and *Fezf2* mutants at P47. For **d-f**, Scale bar: 100 µm. Exact sample genotypes and *P* values are provided in Supplementary Tables 2 and 3.

**Extended Data Fig. 7.**
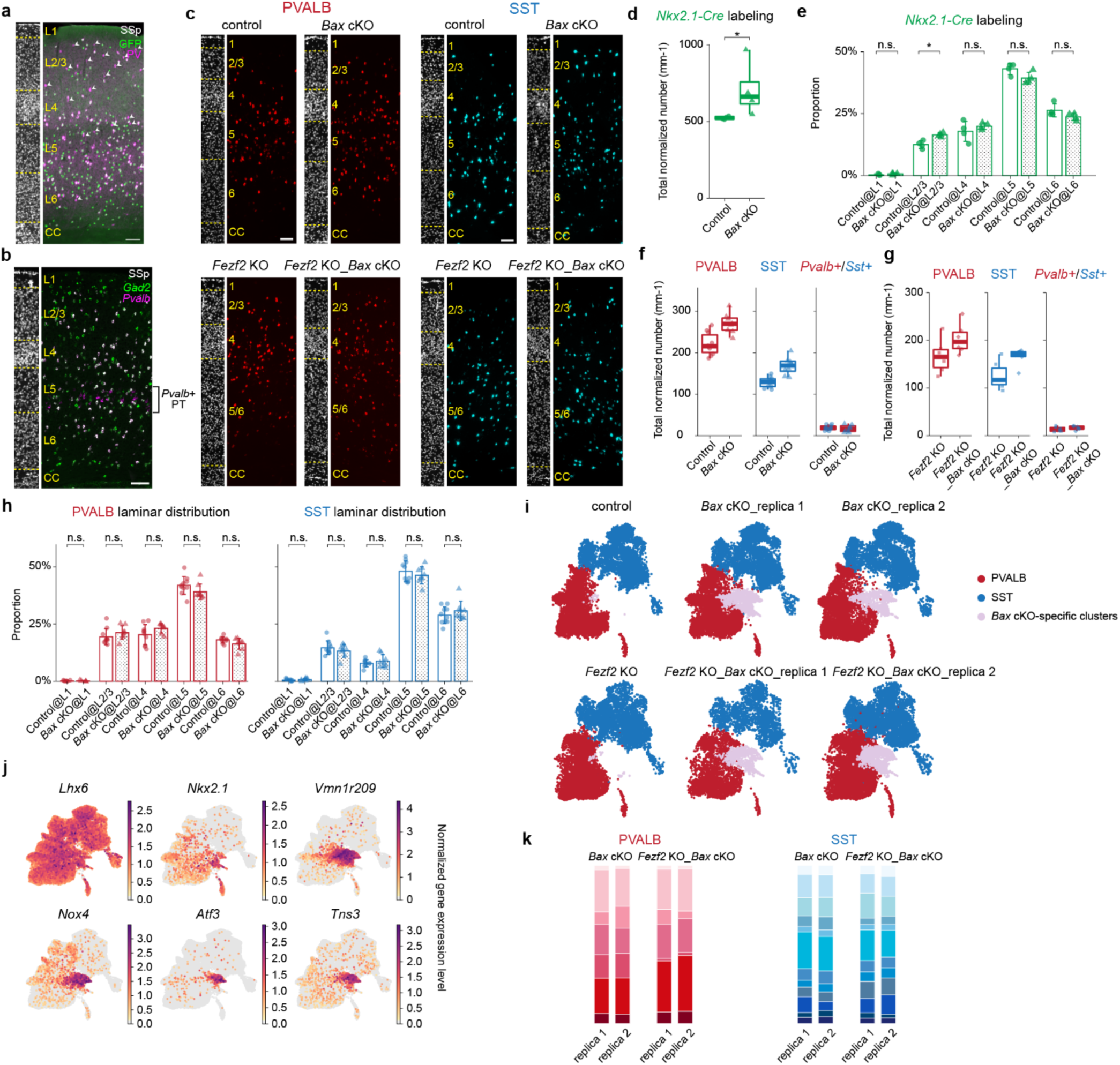
Conditional removal of *Bax* in PVALB and SST interneurons increases their number. **a,** Immunolabeling of PVALB interneurons on *Nkx2.1-Cre* labeled showing the incomplete labeling of superficial interneurons using this driver line. **b,** In SSp region specifically, *Pvalb* mRNA is also expressed in L5b PT neurons (*Gad2*-), which must be excluded from counts. **c,** RNAscope *in situ* hybridization for *Pvalb* and *Sst* mRNA in the SSp region of P28-33 mice with four different genotypes. DAPI counterstaining (left) provides laminar reference. For **a-c**: Scale bar, 100 µm. **d,** Quantification of genetically labeled interneurons by *Nkx2.1-Cre* driver line in control and *Bax* cKO conditions. Sample sizes are presented as n = mice (from X ROIs). control: n=4 (8), n=2316 GFP+ interneurons; *Bax* cKO: n=4 (8), n=3548 GFP+ interneurons. **e,** Laminar distribution of genetically labeled interneurons in control and *Bax* cKO conditions (same dataset as in **d**). Error bars, s.d. **f,** Quantification of PVALB and SST interneurons in control and *Bax* cKO cortices in the SSp region, using RNAscope *in situ* hybridization. L5 PT neurons excluded (*Pvalb*⁺/*Lhx6*⁻). Numbers are normalized to the length of the outskirts of the cortex in millimeters. control: n=5 (10), age P26-28, n=3932 PVALB, n=2280 SST, n=337 *Pvalb+*/*Sst+* interneurons. *Bax* cKO: n=4 (10), age P26-28, n=3470 PVALB, n=2150 SST, n=221 total *Pvalb+*/*Sst+* interneurons. **g,** Quantification of PVALB and SST interneurons in *Fezf2* KO and *Fezf2* KO_*Bax* cKO cortices, based on RNAscope *in situ* hybridization of fresh-frozen sections. *Fezf2* KO: n=4 (6), age P28-33, n=1668 PVALB, n=1257 SST, n=138 *Pvalb+*/*Sst+* interneurons. *Fezf2* KO_*Bax* cKO: n=4 (6), age P28-33, n=2161 PVALB, n=1768 SST, n=175 *Pvalb+*/*Sst+* interneurons. **h,** Laminar distribution of PVALB and SST interneurons in control and *Bax* cKO conditions (same dataset as in **g**). Error bars, s.d. **i,** UMAP plots of P14 snRNA-seq data on interneurons from four different genotypes, showing Bax cKO-specific clusters. **j,** UMAP plots showing normalized expression level of selected genes enriched in *Bax* cKO-specific clusters. **k,** Stacked bar plots showing that the proportion of different PVALB and SST interneurons remains relatively consistent across biological replicates of *Bax* cKO and *Fezf2* KO_*Bax* cKO snRNA-seq datasets. For **e, h,** Wilcoxon rank-sum test, n.s. not significant, p≥0.05; *p<0.05. Exact sample genotypes and *P* values are provided in Supplementary Tables 2 and 3.

**Extended Data Fig. 8.**
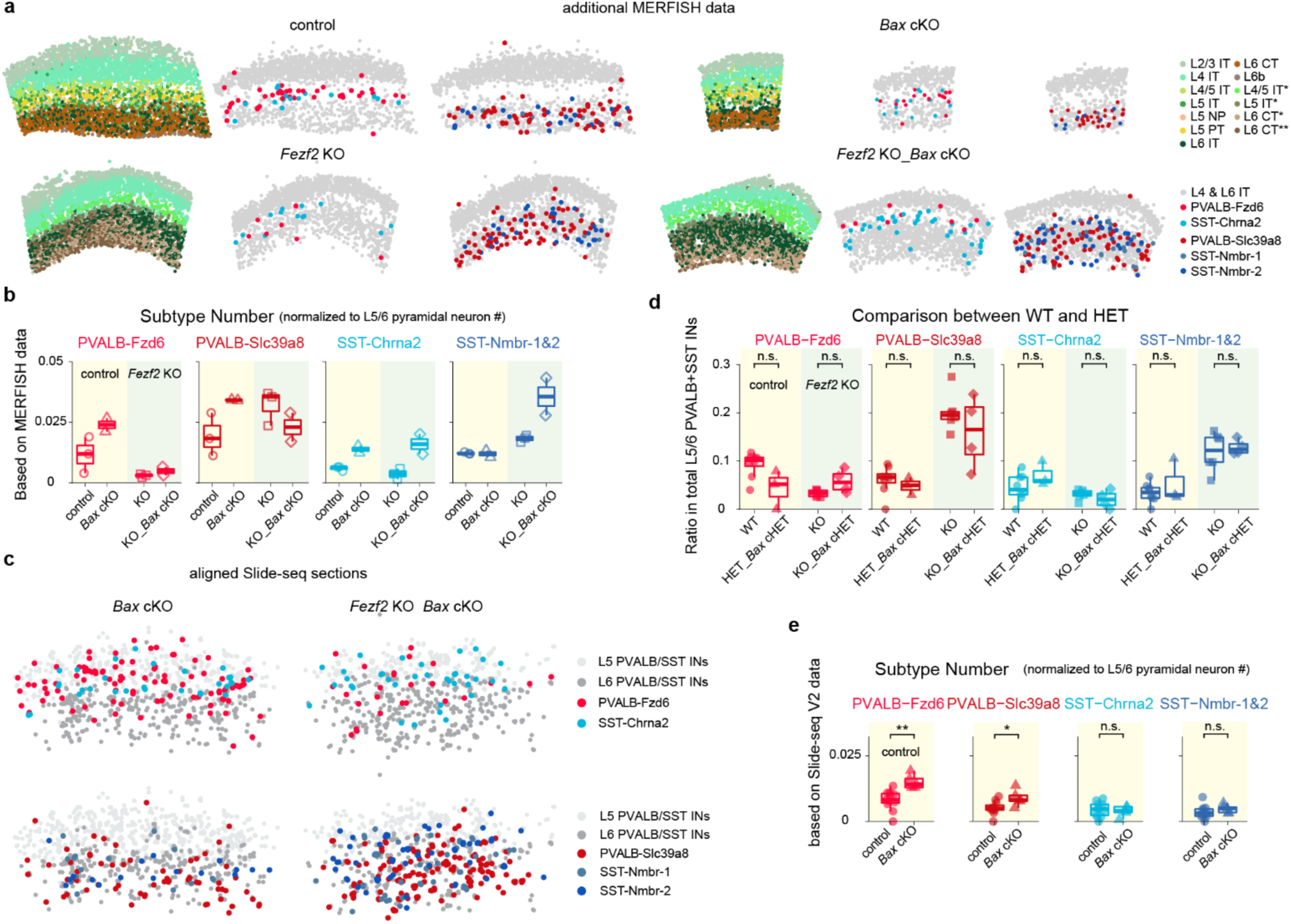
Spatial transcriptomics analysis of *Bax* removal in PVALB and SST interneurons in the *Fezf2* mutant background. **a,** Additional MERFISH data at P14 from different genotypes, complementing Fig. 3d. **b,** Quantification of select subtype numbers in the SSp region, normalized to deep-layer PNs, based on MERFISH data. Note: jthese normalized number is higher than Slide-seq due to higher spatial resolution and cell calling recovery rate offer with this technique. No statistical test due to limited dataset size. **c,** Aggregate Slide-seq maps of coronal sections from *Bax* cKO and *Fezf2* KO_*Bax* cKO mice, highlighting selected PVALB and SST interneuron subtypes identified in L5/6. Sample sizes are presented as n = mice (from X ROIs). *Bax* cKO: n=3 (5); *Fezf2* KO_*Bax* cKO: n=3 (5). Age range: P28-33. **d,** Ratio of selected PVALB and SST interneuron subtypes within total L5/6 PVALB and SST interneurons are consistent between wildtype and heterozygous genotypes in the SSp region, based on Slide-seq data, justifying their pooling for analysis. *Fezf2* WT: n=5 (8); *Fezf2* HET_*Bax* cHET: n=2 (3); *Fezf2* KO: n=5 (6); *Fezf2* KO_*Bax* cHET: n=2 (4). Age range: 4-6 weeks old. These data are included in Fig. 2d. **e,** Numbers of selected PVALB and SST subtypes normalized to L5/6 PN counts in control vs. *Bax* cKO cortices, based on Slide-seq data. control: n=7 (11); *Bax* cKO: n=3 (5). Age range: P28-37. For **d**, **e**, Wilcoxon rank-sum test. n.s. not significant, p≥0.05; *p<0.05; **p< 0.01. Exact *P* values are provided in Supplemental Table 3.

**Extended Data Fig. 9.**
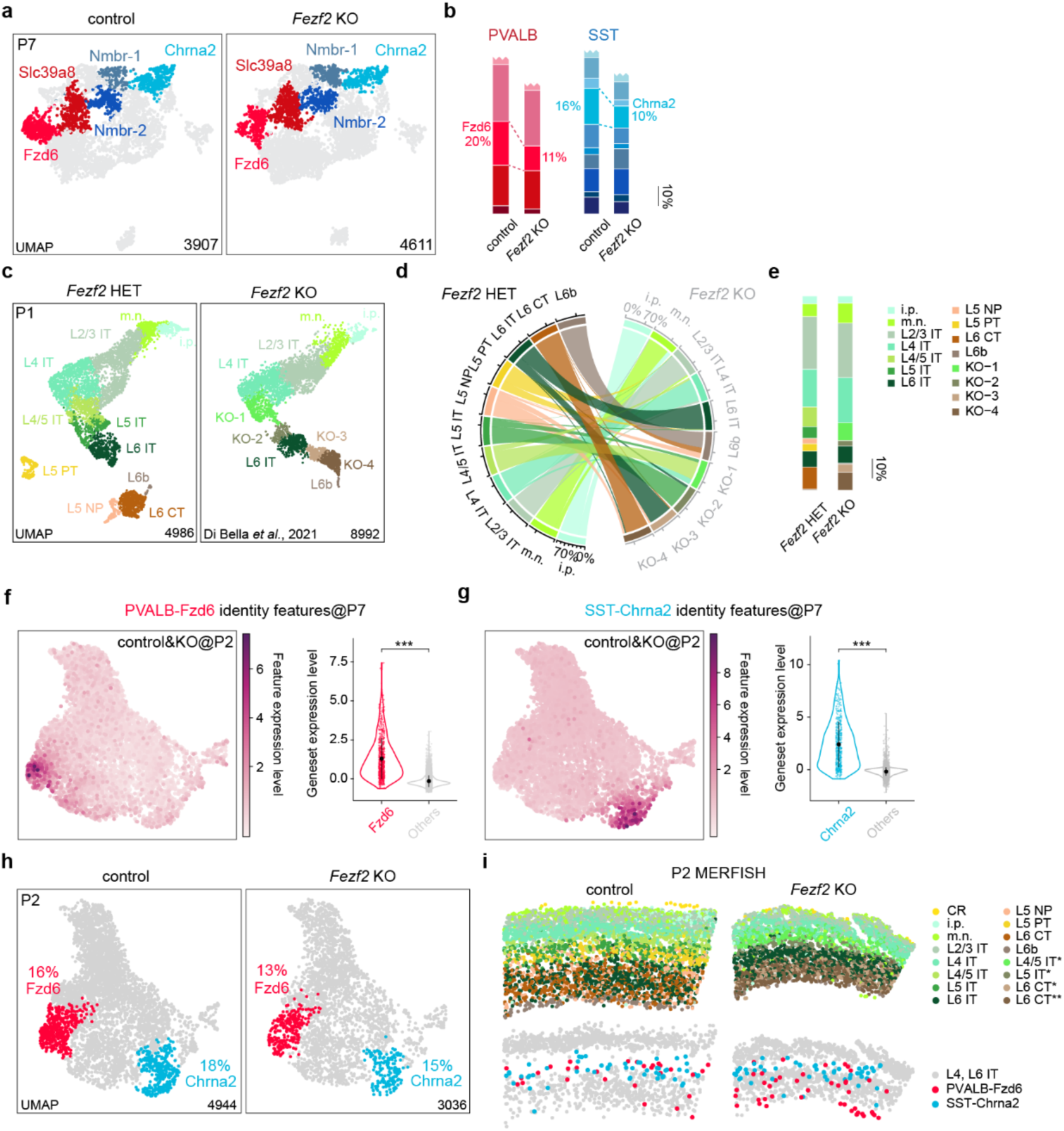
Interneuron subtype changes in *Fezf2* mutants occur after P2. **a,** UMAP visualization of snRNA-seq data of cortical interneurons collected from control and *Fezf2* KO mouse brains at P7, highlighting key deep-layer subtypes. **b,** Stacked bar plots showing the proportion of deep-layer PVALB and SST subtypes in control and *Fezf2* KO brains based on snRNA-seq data. **c,** UMAP visualization of published snRNA-seq data^43^ of excitatory neurons from P1 *Fezf2* HET and *Fezf2* KO mouse cortices. **d,** River plot showing the transcriptomic correspondence between control and KO subtypes. **e,** Stacked bar plots showing PN subtype composition in P1 *Fezf2* HET and KO brains. based on snRNA-seq data. **f-g,** Feature genes identified in P7 snRNA-seq dataset that are selectively expressed in PVALB-Fzd6 and SST-Chrna2 interneuron subtypes are also selectively expressed at P2. (left) UMAP plot showing the expression level of feature genes. (right) Violin plot comparing the expression level of feature genes putative PVALB-Fzd6 and SST-Chrna2 cluster compared to other nuclei in the dataset. Wilcox rank-sum test; ***p<0.001. Exact *P* values are provided in Supplementary Tables 3. **h,** UMAP visualization of snRNA-seq data of cortical interneurons collected from control and *Fezf2* KO mouse brains at P2, with PVALB-Fzd6 and SST-Chrna2 interneuron subtypes highlighted. **i,** MERFISH spatial map of coronal brain sections from the SSp region of control and *Fezf2* KO mice at P2, showing PN and selected interneuron subtypes. CR, Cajal-Retzius cells; i.p., intermediate progenitor; m.n., migrating neurons.

**Extended Data Fig. 10.**
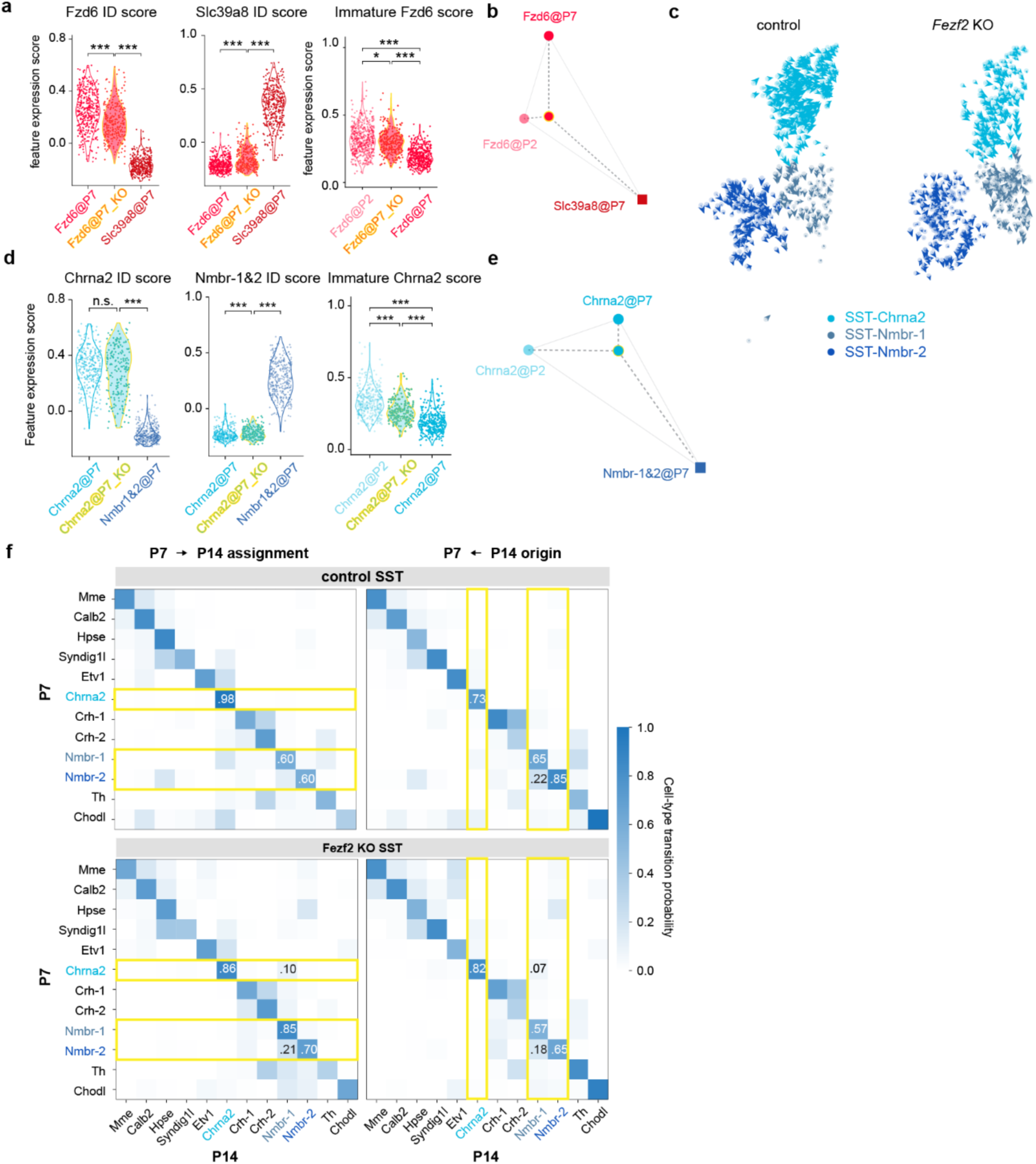
Transcriptomic plasticity of PVALB and SST interneurons in *Fezf2* mutants. **a,** Expression scores for PVALB-Fzd6 (P7), PVALB-Slc39a8 (P7), and immature PVALB-Fzd6 (P2) gene sets (see Methods) in PVALB-Fzd6 interneurons from *Fezf2* mutants, benchmarked against control PVALB-Fzd6 (P7, P2) and PVALB-Slc39a8 (P7). **b,** Triangular Affinity Map of PVALB-Fzd6 interneurons in mutant condition at P7 (see Methods), showing relative transcriptomic similarity to the three reference groups, based on gene set analysis in **a. c,** RNA velocity analysis of snRNA-seq data on SST interneurons from control and *Fezf2* KO cortices at P7, showing gene expression dynamics of three SST subtypes. **d-e,** gene set analysis and Triangular Affinity Map, as in **a-b**, for SST-Chrna2 interneurons in *Fezf2* mutants at P7, benchmarked against control SST-Chrna2 (P7, P2) and SST-Nmbr-1&2 (P7). **f,** Optimal transport analysis mapping the developmental trajectories of SST subtypes from P7 to P14 in control and *Fezf2* KO conditions. Descendent analysis (left) shows predictive assignment from P7 to P14 subtypes; ancestor analysis (right) shows predicted P7 origins of P14 subtypes. For **a**, **d**,Wilcoxon rank-sum test, *p<0.05; ***p<0.001. Exact *P* values are provided in Supplementary Tables 3.

**Extended Data Fig. 11.**
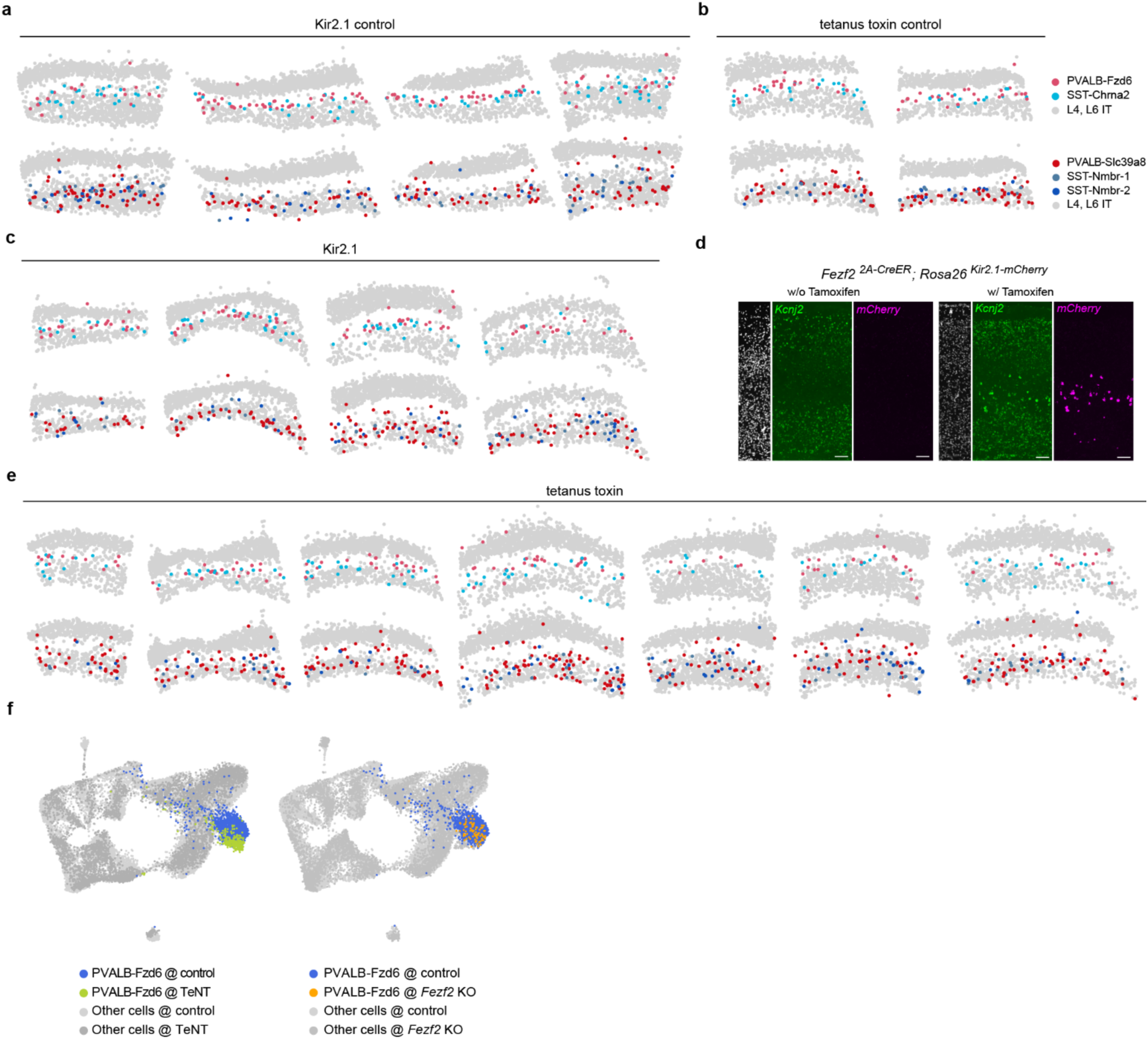
Additional Spatial data and transcriptomic validation of PN manipulation experiments. **a-c, e,** Additional MERFISH spatial maps of control, Kir2.1, and TeNT conditions, supplementing Fig. 5a and contributing to quantification in Fig. 5b. **d,** RNAscope in situ hybridization confirms the induced expression of *Kcnj2*, the gene encodes for Kir2.1, in *Fezf2*+ L5b PNs. **f,** UMAP visualization of integrated snRNA-seq datasets from control, *Fezf2* KO (dataset as in Fig. 2a), and TeNT conditions. The analysis compares the transcriptomic state of PVALB-Fzd6 interneurons in the TeNT condition (left) and the *Fezf2* KO condition (right) relative to combined controls, revealing a more pronounced transcriptomic shift in the remaining cells of the TeNT condition. Exact sample genotypes are provided in Supplementary Tables 2.

**Extended Data Fig. 12.**
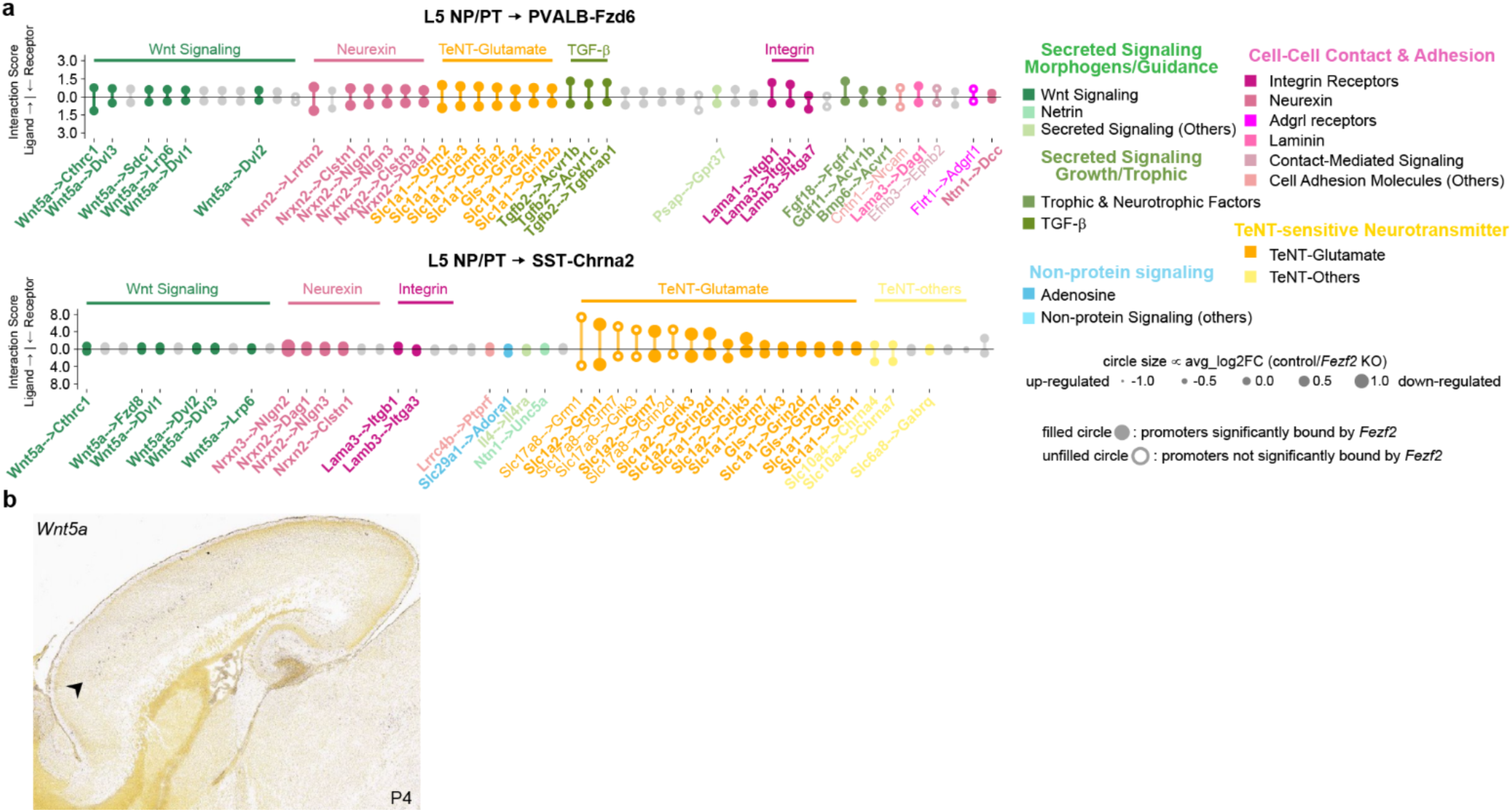
Additional candidate ligand–receptor interactions and *Wnt5a* expression. **a,** Lollipop diagram, as in Fig. 5c, showing the 51st-100^th^ ranked candidate ligand/receptor pairs from the bioinformatic screen. See **Supplemental Tables 4 and 5** for a complete list of the top 100 ligand-receptor pairs.**b,** Representative in situ hybridization image for *Wnt5a* at P4, showing specific expression in L5b. Image from Allen developing mouse brain atlas: in situ hybridization (ISH) data^34^ (https://developingmouse.brain-map.org/experiment/show/100101325).

## Methods

### Mice

All experimental procedures were approved by the Harvard Medical School Institutional Care and Use Committee and were performed in compliance with the Guide for Animal Care and Use of Laboratory Animals. Mice were housed in a temperature-controlled and humidity-controlled facility and were maintained on a 12–12 h dark-light cycle. All experiments were performed on animals of both sexes. Whenever possible, mice of both sexes are used in experiments. Experiments were not blinded because mice and treatments were easily identifiable as experiments were performed. Sample sizes were not predetermined.

Mouse lines were used in this study include *Fezf2*^−/−^ (Ref ^44^), *Rorb^Cre^* mice (JAX #023536), *Tcerg1l^CreER^* mice (JAX #034000), *Pvalb^FlpO^* (JAX #022730), , *Dlx5/6-Cre* (JAX #008199), *Sst^Cre^* (JAX #018973), *Sst^FlpO^* (JAX #031629), *Chrna2-Cre* (Ref ^13^), *Pdyn^Cre^* (JAX #027958), *Npy^FlpO^* (JAX #030211), *Crhr2^Cre^* (JAX #033728), *Hpse^Cre^* (JAX #037334), *Nkx2.1-Cre* (JAX #008661), *Fezf2^2A-CreER^* (JAX #036296), *Emx1^IRES-Cre^*(JAX #005628), *Fezf2^2A-FLpO^* (JAX #036297), *Bax^fl^*(MGI ID:3589203), *Rosa26^Kir2.1−mCherry^* (MGI:6191641), *RC::PFtox* (MGI:4412286), *Rosa26^FSF-LSL-tdTomato^*(*Ai65)* (JAX #021875), *Rosa26^LSL-tdTomato^* (*Ai14*) (JAX #007914), *Rosa26^CAG-Sun1/sfGFP^ (INTACT)* (JAX #030952).

### snRNA-seq

#### Nuclei isolation and snRNA-seq library construction

Cortices were dissected from mice of specific genotypes and ages in ice-cold Homogenization Buffer (HB: 0.25 M sucrose, 25 mM KCl, 5 mM MgCl_2_, 20 mM Tricine-KOH, 1 mM DTT, 0.15 mM spermine, and 0.5 mM spermidine). The dissected brain tissue was transferred to a 2-mL Dounce tissue grinder containing HB supplemented with 0.15-0.25% IGEPAL-CA630 and 0.2 U/µl RNasin. Tissue homogenization was performed with 8-10 strokes of pestle A, followed by a 5-min incubation on ice, and then 9-10 strokes of pestle B. The homogenate was filtered through a 30 μm strainer into 15 ml tube and centrifuged at 500 g for 7 min at 4 °C (swinging-bucket rotor). The resulting pellet was resuspended in 1X PBS with 1% BSA and 0.2 U/µl RNasin, then passed through a 40 μm filter. Based on genetic, viral, and nuclear dye labeling (see Supplemental Methods), nuclei were sorted using a Sony MA900 cell sorter, collected, concentrated as needed, and counted with a hemocytometer (INCYTO C-Chip). snRNA-seq libraries were prepared using the Chromium Single Cell 3’ Kit (v3.1 or v4, 10X Genomics) according to the manufacturer’s protocol. Pooled libraries were sequenced on Illumina NovaSeq 6000 instruments.

#### snRNA-seq data processing

Raw sequencing data was processed using CellRanger (v7.0.0, default parameter, 10X Genomics) to generate gene expression matrices by aligning reads to the mouse reference genome (mm10). The resulting matrices were imported into Python (v3.9.13) as AnnData objects (anndata v0.8.0), and all subsequent analyses were performed using Scanpy (v1.9.1). For quality control, we filtered out genes expressed in fewer than 3 cells, unless otherwise noted. We then calculated a doublet score for each nucleus using Scrublet (v0.2.3) ^45^ and only retained cells that met all the following criteria: n_counts < 15,000, 200 < n_genes < 4,000, ratio_of_mitochondria_genes <5% (genes prefixed “mt-”), ratio_of_ribosome_genes <5% (genes prefixed “Rpl/Rps”), and Scrublet score <0.25. Raw counts of filtered data were normalized to 10,000 counts per nucleus (pp.normalize_total) and log-transformed (pp.log1p). Highly variable genes (HVGs) were selected from raw counts using Pearson residuals (experimental.pp.highly_variable_genes; *n* = 5,000). HVG residuals were scaled followed with calculating the Principal Component Analysis (PCA) cell embeddings using scikit-learn v.0.24.0 (StandardScaler: with_mean=False; TruncatedSVD). Unsupervised graph-based Leiden clustering is performed based on PCA embedding (pp.neighbors; tl.leiden). For targeted subclustering, we used the piaso.tl.leiden_local function from PIASO, which re-runs HVG selection, PCA and Leiden clustering on the selected subset, while preserving clustering assignments for the remaining cells. Cell type annotation was then achieved by identifying marker genes for each cluster with COSG^46^, and cross-reference to known markers in the literature. For visualization of multiple snRNA-seq datasets across different conditions, the piaso.pl.plot_embeddings_split function from PIASO was used to align cell embedding across datasets and to scale gene expression or cell metrics for consistency.

#### Source and processing of public scRNA-seq datasets

We downloaded and processed several public scRNA-seq and snRNA-seq datasets. They include 10X Genomics datasets: P28 mouse cortex interneuron snRNA-seq (GEO: GSE164570)^11^, P14 control and *Fezf2* KO mouse S1 cortex snRNA-seq datasets (GEO: GSE158096)^26^, and P1 *Fezf2* Het and *Fezf2* KO mouse cortex scRNA-seq datasets (GEO: GSE153164)^43^. For each, FASTQ files were reprocessed with CellRanger (v7.0.0) with default parameters as described above. Additionally, processed and annotated SMART-Seq V4 scRNA-seq datasets were downloaded from previous publications^12,47^ and re-annotated to ensure consistent cell type labels.

#### Individual cell-based differential transcriptomic analysis across conditions (Emergene)

To assess transcriptomic differences across conditions at single-cell resolution, we developed Emergene, a clustering-agnostic framework designed to reduce cluster-size bias common to differentially expressed gene (DEG) workflows. For each gene, Emergene builds two diffusion maps of its expression vector on a shared low-dimensional embedding: (i) a local map obtained by diffusing to the *k* nearest neighbors (kNN), and (ii) a null map obtained by diffusing to *k* randomly chosen cells. Deviation from the original expression vector is quantified by cosine similarity (as in COSG⁴⁰). Genes with localized expression are selected based on the difference between the local and null deviations. Using a similar diffusion strategy on a shared cell embedding space, the algorithm then identifies a key set of genes whose local co-expression patterns are altered among experimental conditions. Emergene aggregates the localized and condition-specific genes into a combinatorial gene set and computes a normalized enrichment score per cell. Statistical significance is assessed via a permutation test adapted from scDRS: for each tested set, Emergene draws 10,000 control gene sets matched on size, mean expression, and variance to form an empirical null. Cell-level *P* values are derived from the distribution of normalized scores across all cells and matched controls. For each cluster (subtype), Emergene reports a summary significance using a gliding threshold on the distribution of cell-level *P* values (median, 50th percentile, by default).

#### Feature gene set identification and expression scoring

The identity (ID) feature gene set of a specific interneuron subtype was comprised of the top 200 marker genes identified by COSG that were differentially expressed between the two PVALB or SST subtypes. The immature feature genes were the top 200 DE genes identified by COSG that expressed higher at P2 than at P7 for specific interneuron subtypes. DE genes between control and *Fezf2* KO conditions were the top 200 DE genes identified by COSG across conditions for specific interneuron subtypes. The enrichment of each of these gene sets in individual cells was calculated as a feature score with scanpy.tl.score_genes (Scanpy v1.9.1) using default parameters.

#### Triangular Affinity Map for comparative visualization of gene expression similarity

To generate a visual representation of transcriptomic similarity between one target cell type and three anchor cell types, we employed the Triangular Affinity Map (TAMap) based on gene set expression scores. We first applied min-max normalization to adjust the mean gene set expression scores across various cell type comparisons to a uniform scale, ensuring consistency in the similarity metrics. We subsequently calculated the sizes of three internal angles, each spanned by the target cell type and one of the anchor cell types. These angles represent the similarities between two anchor cell types—such that their sum equaled 360 degrees. Additionally, we computed the edge lengths connecting the target cell type with each anchor cell type based on the similarities between the target cell type and each anchor cell type. For a 2D visualization in a Cartesian coordinate system, the target cell type was positioned at the origin, and one anchor cell type along the positive y-axis. Matplotlib v3.5.2 was utilized to generate the final visual representation.

#### RNA velocity

To predict how the gene expression pattern of each cell type will change developmentally, we employed scVelo v0.2.4^32^ to infer the RNA velocity from the ratio of spliced and unspliced reads. STAR 2.7.10a was used to quantify the counts of spliced and unspliced reads in individual cells from the P7 *Fezf2* HET and P7 *Fezf2* KO snRNA-seq FASTQ files. Next, we mapped the information of annotated cell types and UMAP coordinates based on cell barcodes. RNA velocities were computed using the scvelo.tl.recover_dynamics, scvelo.tl.velocity(mode=’dynamical’) and scvelo.tl.velocity_graph() functions, and visualized based on UMAP coordinates with the scvelo.pl.velocity_embedding() function.

#### Marker <u>G</u>ene-guided <u>D</u>imensionality <u>R</u>eduction (GDR)

GDR is developed for dimensionality reduction and data integration for snRNA-seq across ages or conditions. It first identifies marker genes across clusters using COSG^46^. GDR calculates expression scores for these marker gene sets across datasets, projecting cells into a shared low-dimensional gene expression space that reflects biological identity in each dimension. In this space, cells with similar identities or gene expression patterns are located closely together. The GDR code is available in the PIASO GitHub repository (piaso.tl.runGDR).

#### Optimal transport

Cell fate transition probabilities between P7 and P14 were quantified for PVALB and SST lineages separately in control and *Fezf2* KO conditions using an optimal transport framework (Moscot v0.4.3^33^). For each comparison, a cost matrix was defined based on distances within a 30-nearest neighbors graph built on the GDR cell embedding. The moscot.problems.time model was then solved with default settings (epsilon=1e-3, scale_cost=“mean”, max_iterations=1e7) to obtain a probabilistic coupling between the two time points. This coupling yields forward transition probabilities (descendant analysis/cell fate assignment) and backward probabilities (ancestor analysis/cell fate origin) between interneuron subtypes.

#### Ligand/receptor bioinformatics search

To nominate the ligand-receptor (L-R) pairs that mediate the interactions between L5 PT/NP neurons and L5b PVALB and SST interneuron (IN) subtypes, we analyzed P4 and P8 mouse cortex scRNA-seq datasets released by the BRAIN Initiative Cell Atlas Network (BICAN Rapid Release Inventory; https://www.portal.brain-bican.org/rapid-release). These datasets were preprocessed and annotated using the same pipeline described above. Candidate L–R pairs were drawn from the CellChat^48^ database, then manually curated and supplemented with additional relevant entries. We developed SCALAR (<u>S</u>ingle <u>C</u>ell <u>A</u>nalysis of <u>L</u>igand <u>A</u>nd <u>R</u>eceptor), implemented in the PIASO toolkit, to identify subtype-specific interactions. SCALAR was based on COSG^46^. It first calculates the expression specificity scores for ligands across all PN subtypes and for receptors across all IN subtypes. These scores were normalized with the cosg.iqrLogNormalize function to enable cross-subtype comparison. For each PN-IN pair, SCALAR then calculated an interaction score for individual L-R pair in the curated database. The significance of each L-R pair is assessed by a permutation test, where SCALAR randomly sampled 1,000 L-R pairs (matched on mean and variance of L-R expression across all the cells with the L-R to be tested) to form a null distribution. Within each PN-IN pair, the False Discovery Rate (FDR) via Benjamini-Hochberg procedure was then calculated using the statsmodels.stats.multitest.multipletests function in the statsmodels v0.14.4 package, and only ligand-receptor pairs with FDR <0.2 are retained. The SCALAR code is available in the PIASO GitHub repository (piaso.tl.runSCALAR). The L-R pairs were ranked by the interaction scores, and top L-R pairs were visualized via piaso.pl.plotLigandReceptorLollipop function. To further prioritize the L-R pairs, we employed MAST^49^ to identify ligands which were differentially expressed in *Fezf2* KO versus control at P7, and utilized the public *Fezf2* ChIP-seq data^35^ to identify which ligands could be directly bound by *Fezf2* in their promoter regions.

### MERFISH

#### Selection of probe set

To design the MERFISH gene panel that captures the transcriptomic heterogeneity across cell types, developmental stages and genotypes, we utilized multiple snRNA-seq datasets collected across a wide age range. Top marker genes identified by COSG were combined with known marker genes from literature to generate an initial gene panel for late postnatal ages. Additional genes were added and tested on scRNA-seq collected from younger age to ensure comprehensive cell type identification across age. RNA targets were selected based on maximizing unique probe sites per gene for high detection rate while minimizing combinatorial optical crowding for MERFISH imaging. A fluorescently labeled oligonucleotide library probing for 293 combinatorial genes and 4 sequential genes, including GFP was selected. The resulting readout and encoding probes were manufactured by Vizgen Inc. The full list of probes is available in Supplemental Table 1.

#### MERFISH sample processing

Whole intact brain samples were collected in RNase-sterile conditions, flash-frozen in liquid nitrogen, and moved to 5 mL Eppendorf tubes and stored at −80 °C for tissue microarray (TMA) construction and MERFISH. First, frozen cortical tissues were collected using 2 mm or 3mm disposable biopsy punch needles from specific brain regions. The tissue punches were then trimmed uniformly with a sterile razor blade, and oriented laterally and embedded within a pre-formed scaffold of Optimal Cutting Temperature media. On average, six sample punches were assembled into each TMA. All samples were prepared in RNase-sterile conditions for MERFISH imaging according to the procedure described in a previous publication^50^ and using select additional kits and instruments offered through Vizgen Inc. Briefly, the TMA samples were cryosectioned at 10 μm using a cryostat (Leica) at -20°C, and mounted and melted onto fluorescent microsphere-coated, functionalized coverslips, fixed with 4% PFA in 1X PBS, and permeabilized overnight in 70% ethanol. TMA sections were stained using the Cell Boundary Stain Kit (Vizgen, PN 10400009). Following a 1-hour room temperature blocking step in Cell Boundary Block Buffer containing 40 U/µl murine RNase inhibitor, samples were incubated in the primary and secondary antibody cocktails for 1 hour each, with interspersed 1X PBS washes. Primary antibodies against specific proprietary cell membrane proteins and oligo-conjugated secondary antibodies were diluted in Cell Boundary Block Buffer containing 40 U/µl murine RNase inhibitor, with dilution factors of 1:100 and 3:100, respectively. Antibody labeling was fixed again with 4% PFA for 15 minutes. TMA sections were then hybridized with 70 μL of 297-gene probe library solution for 36-48 hrs in a humidified 37 °C incubator with a 2×2 cm square of Parafilm layered onto the surface to prevent evaporation. Samples were embedded in a polyacrylamide gel by incubating the samples in freshly prepared polyacrylamide gel solution (40% 19:1 acrylamide/bis-acrylamide solution, 5M NaCl, 1M Tris PH8, and nuclease-free water in a dilution of 1:3:3:39; gel solution combined with catalysts, 10% w/v ammonium persulfate solution and NNN’Tetramethyl-ethylindiamin in a dilution of 2000:10:1). To achieve this, the coverslip containing TMA samples were inverted onto the polyacrylamide solution aliquoted on the surface of a Gel-Slick treated, 2×3 inch microslide. Non-targeted molecules were cleared from the gel-embedded sample within a detergent mixture (20X saline-sodium citrate, 10% sodium dodecyl sulfate solution, 25% Triton-X 100 solution, and nuclease-free water in a 5:10:1:34 ratio) supplemented with 0.8 U/µl Proteinase K (NEB P8107S) for 48-72 hours in a humidified 37 °C incubator.

#### MERFISH imaging and post-imaging processing

Samples were further stained with DAPI and PolyT Staining Reagent (Vizgen, PN 20300021), and imaged using MERSCOPE (Vizgen) with MERSCOPE 300-Gene Imaging Cartridges (Vizgen, PN 20300017). Illumination intensities and exposure times were kept the same in every dataset, capturing images of the whole TMA with both a 10× NA 0.25 and 60× NA 1.4 objective, at 7 z-positions per x–y location, separated by 1.5 µm. MERSCOPE post-imaging analysis used MERlin^51^ to decode positions and copy numbers of target RNA species into count matrices, and Cellpose^52^ to generate cell boundary masks (software version 233.230615.567, segmentation parameter: cell boundary 1).

#### Cell type identification for MERFISH data

After quality control, four major classes of cell types were identified based on marker gene expression: excitatory neurons; two groups of interneurons (PVALB/SST and VIP/SNCG/LAMP5 groups); and non-neuronal cells. Each class was separately annotated using the GDR. In brief, GDR projected cells from snRNA-seq and MERFISH into a shared low-dimensional gene expression space that reflects biological identity in each dimension, placing cells with similar identities in closer proximity. To mitigate cross-modality technical effects, we used the pp.harmony_integrate function from Scanpy v1.9.1 with default parameters to run the Harmony^53^ integration procedure. Following this, Support Vector Machine (SVM) with the radial basis function kernel was used to predict the cell types of cells in the MERFISH data based on the cell embeddings from both MERFISH and snRNA-seq data. SVM was implemented through the svm.SVC function from scikit-learn v.0.24.0, set to kernel=’rbf’, and class_weight=’balanced’.

#### Analysis of Allen MERFISH dat

The MERISH spatial transcriptomics data of a single adult mouse brain (Allen MERFISH data) with a 500 gene panel was published in Yao *et al.*, 2023^12^. The processed and spatially aligned data with raw counts was downloaded from Allen Brain Cell Atlas (C57BL6J-638850-raw.h5ad; https://alleninstitute.github.io/abc_atlas_access/descriptions/MERFISH-C57BL6J-638850.html). We subset the MOp, MOs, SSp and VISp regions based on the “parcellation_structure” annotation as the data was registered to the Allen CCFv3 (Common Coordinate Framework). The dataset was also further divided into four major cell type groups and annotated separately using snRNA-seq or scRNA-seq reference with GDR as described above. Specifically, 10x Genomics v3 snRNA-seq datasets was used as reference for interneurons. SMART-Seq v4 scRNA-seq datasets were used as references for excitatory neuron and non-neuronal cells. QC and cell type annotation of these reference datasets were performed as described above.

### Slide-seq

#### Library generation and sequencing

Slide-seq pucks (round, 3 mm in diameter) were generated as described previously^54^. 10 µm-thick coronal sections were obtained from flash-frozen brain samples using cryostat (Leica) and used for generating Slide-seq library immediately, following published Slide-seqV2 protocol^42,55^. Libraries were pooled and sequenced on NovaSeq 6000 flow cells (Illumina). One published Slide-seq data on mouse cortex SSp region was included in this study, puck 200306_02, can be downloaded at https://singlecell.broadinstitute.org/single_cell/study/SCP815/highly-sensitive-spatial-transcriptomics-at-near-cellular-resolution-with-slide-seqv2.

#### Slide-seq data pre-processing

The sequenced reads were aligned to GRCm39.103 reference and processed using the Slide-seq tools pipeline (https://github.com/MacoskoLab/slideseq-tools; v.0.2) to generate the gene count matrix and match the bead barcode between array and sequenced reads. The spatial barcode recovery step was further optimized with customized algorithm.

#### Cell type mapping using RCTD

We used RCTD^56^ (now available in the R package spacexr v2.2.1) to map cell types in Slide-seq data based on reference scRNA-seq data. The reference dataset was processed by first retaining only the genes detected in both the Slide-seq data and the reference scRNA-seq data. We then used the R version of COSG (v0.9.0) with the following parameters: mu=100, remove_lowly_expressed=TRUE, expressed_pct=0.1, to select the union set of the top 100 marker genes for each cell type in the annotated scRNA-seq reference. This processed scRNA-seq dataset was then used as input for RCTD. The extraction of distinctive features for each cell type in the reference dataset increased the accuracy of cell type decomposition in Slide-seq data. From the RCTD output, we retained beads classified as ‘singlet’ or ‘doublet_certain’ and excluded the rest.

#### Inference of interneuron laminar locations in MERFISH and Slide-seq data

The resident cortical layer for each interneuron was assigned based on the identity of its five nearest excitatory neuron neighbors. Specially, the KDTree function from scikit-learn v.0.24.0 with leaf_size=6 and k=5 was used for this purpose.

### Statistics

Most of the statistical tests in this study employed two-sided non-parametric tests due to the small sample sizes (n<25) and the non-normal distribution of single-cell expression data. Boxplots display the median, the 25^th^ percentile, the 75^th^ percentile, with two whiskers that extend to 1.5 times the interquartile range. A summary of statistical test results is provided in Supplementary Table 2.

## Data availability

The data have been deposited at GEO under accession number GSE272706 and at the Single Cell Portal: https://singlecell.broadinstitute.org/single_cell/study/SCP2716/pyramidal-neurons-control-the-number-and-distribution-of-cortical-interneuron-subtypes.

## Code availability

Emergene is available via https://github.com/genecell/Emergene, and PIASO is available via https://github.com/genecell/PIASO.

## Acknowledgements

We thank P. Arlotta and S. Lodato for providing insightful discussions and suggestions on the manuscript. We thank P. Arlotta for providing the *Fezf2* KO and *Tcerg1l^CreER^* mouse line; P. Keeley for providing the *Bax^fl^* mouse line; K. Kullander for providing the *Chrna2-Cre* mouse line; D. Ginty for providing the *Rorb^Cre^* mouse line; D. A. Stafford for providing the *Hpse^Cre^* mouse line. This work was supported by grants from the National Institutes of Health (NIH) R01NS081297 and R37MH071679 to G.F., the William Randolph Hearst Fund (FY20) to S.J.W., F32 fellowship from National Institute of Mental Health (NIMH) F32MH125464 to S.J.W, grant from NIMH 1U01MH130962 to C.M. We thank the Neurobiology Department and the Neurobiology Imaging Facility for consultation and instrument availability that supported this work. Finally, we thank R. Raichur for her assistance on pre-processing the Slide-seq data, and S. Du for additional technical assistance.

## Author Contributions

Conceptualization: S.J.W., G.F. Investigation: S.J.W., S.P.Y., C.M., Y.Q., S.H., D.S. Formal Analysis: M.D., S.J.W. Resources: V.K., G.J.M., E.Z.M., F.C., S.L.F., Q.X., J.A.S., D.J.D. Writing – Original Draft: S.J.W., M.D., C.M., G.F. Writing – Review & Editing: S.L.F., J.T., B.C. Supervision: G.F.

