## Supplementary file list for "Pyramidal neurons proportionately alter the identity and survival of specific cortical interneuron subtypes"

Supplementary Methods: Additional methodology description for FACS procedure, immunostaining, stereotaxic AAV injection, RO AAV injection, Tamoxifen administration, RNAscope *in situ* hybridization, Image Acquisition.

Supplementary Table 1: List of MERFISH probe set

Supplementary Table 2: Sample genotypes

Supplementary Table 3: Statistical test and *P* values

Supplementary Table 4: Top 100 ligand-receptor pairs for PVALB-Fzd6

Supplementary Table 4: Top 100 ligand-receptor pairs for SST-Chrna2
