## Supplemental Methods for "Pyramidal neurons proportionately alter the identity and survival of specific cortical interneuron subtypes"

**Supplementary Methods**

**Fluorescence activated cell sorting of genetically labeled cortical interneurons**

Nuclei were first gated by DRAQ5+ signal, on the peak representing single nuclei. Nuclei expressing fluorescent proteins were then selected as GFP+ single nuclei. An example of the gating strategy is shown in Fig. S1.

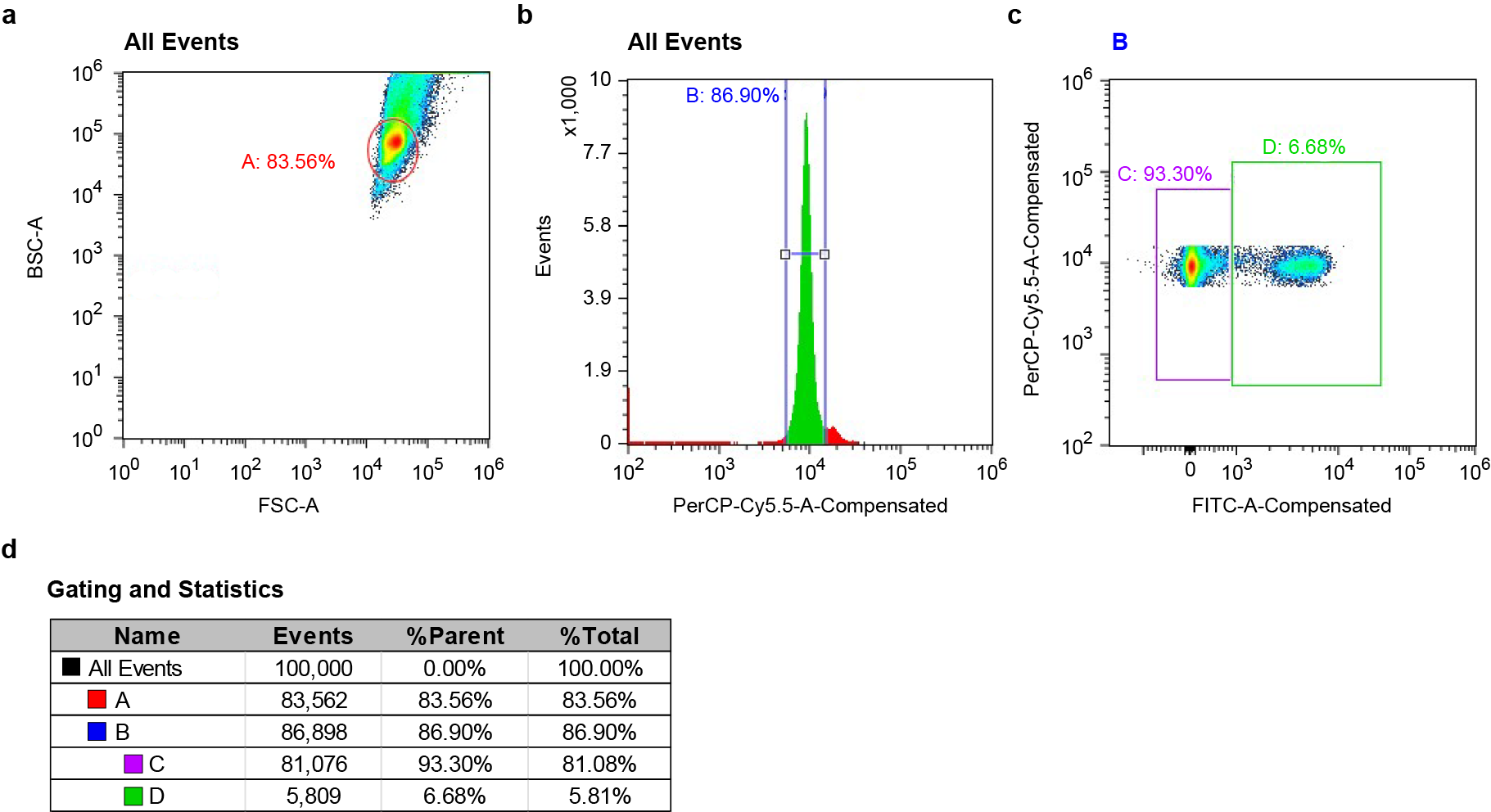

Fig. S1 Example of FACS gating strategy

This example shows FACS of genetically labeled interneurons from a P14 *Nkx2.1-Cre;Bax^fl/+^;Rosa26^CAG-Sun1/sfGFP^* mouse. **a,** Density plots created by Cell Sorter Software (Sony) showing backward scatter area (BSC-A) versus forward scatter area (FSC-A) distributions for the majority of nuclei, circled in A. **b,** Histogram of DRAQ5 signal. A gate (B) was created around the peak representing single nuclei. **c,** The level of GFP signal was used to gate for genetically labeled interneurons (gate D). **d,** The Gating hierarchy and sorting statistics is provided in this table.

**Immunohistochemistry**

Immunohistochemistry experiments were performed to amplify the fluorescent signal of genetic or viral labeled interneurons, or to label PVALB interneurons. For all histological experiments, mice were deeply anesthetized with sodium pentobarbital (Euthasol) by intraperitoneal injection and transcardially perfused with 1X PBS followed by 4% paraformaldehyde (PFA) in 1X PBS. Brains were dissected out and post-fixed overnight at 4°C. Brain slices ranging from 40-150 µm were obtained from sectioning using a vibratome (Leica VT 1200S) or using a microtome (Leica SM2010 R) after cryopreservation. Brain slices may be stored in Storage Buffer at -80 °C before further processing. Free-floating brain sections were incubated in primary antibodies diluted in antibody incubation solution (5% normal donkey serum, 0.25% Triton X-100 in 1X PBS) in coldroom overnight or up to three days. Secondary antibodies were diluted in antibody incubation solution at RT for 1-3 hrs, or in coldroom overnight. Primary antibodies used targeted: GFP (Sicgen AB0020-200), DsRed (Clontech #632496), PV (Swant PV27). Secondary antibodies used include Alexa 488 donkey anti-goat (Thermo Fisher Scientific A-11055) and Alexa 594 donkey anti-rabbit (Thermo Fisher Scientific A-21207).

**Stereotaxic AAV injection**

AAV-PHP.eB-hDlx-DIO-ChR2-mCherry were produced by Neurotools (University of Carolina at Chapel Hill) from user supplied plasmid preparation. To label SST-Hpse and SST-Crhr2 in control and *Fezf2* KO conditions, 100 nL of this AAV (diluted to titer: E+12 vg/ml) is injected in S1 region using a Nanoject III at 6-8 weeks age. The labeling pattern was examined approximately two weeks after injection.

**Retro-orbital injection of in-house prepared AAV**

To label interneurons in *Emx1^Cre^;Fezf2^2A-FlpO^;RC::PFtox* mice, we performed retro-orbital injection of an AAV virus driven by Dlx enhancer at P14. This labeling enabled the subsequent FACS enrichment of interneurons for snRNA-seq expeirments.
Mice were anesthetized with isoflurane (1–3% in oxygen at 0.5 L/min) delivered through an anesthesia induction chamber and maintained using a nose cone system. Depth of anesthesia was assessed by bilateral toe pinch prior to proceeding. Viral delivery using CN3798 (Php.eb serotype) was performed by retro-orbital injection at P14. Each animal received 4.0 × 10^11^ viral genomes in a total volume of 40 μL, administered into the retro-orbital sinus using a 0.3 mL insulin syringe (BD Biosciences, #32470). In-house prepared rAAVs (PHP.eB serotype) were produced as previously described (PMID: 30626963). AAVpro HEK293 cells were transfected using Transporter 5 with three plasmids: the pAAV transfer vector containing the recombinant genome of interest, pUCmini-iCAP-PHP.eb (Addgene 103005) encoding the AAV Php.eb rep/cap protein, and pHelper encoding adenoviral helper proteins. Virions were harvested 72 h post-transfection and purified by iodixanol gradient ultracentrifugation. Viral titers (genome copy number/mL) for in-house prepared CN3798 (Php.eb serotype) was determined by droplet digital PCR (ddPCR) using primers directed against the AAV ITR sequence.  A calibration standard consisting of a virus preparation with an established titer was processed in parallel to normalize copy number estimates.

AiP13798 - pAAV-DLX2.0-minBG-SYFP2-P2A-mScarlet-10aa-H2B-WPRE3-BGHpA (Alias: CN3798) was a gift from The Allen Institute for Brain Science & Jonathan Ting (Addgene plasmid # 229935, RRID:Addgene_229935)

**Tamoxifen administration**

To label PVALB-Fzd6, a varying dose of tamoxifen, ranging from a single dose of 0.5 mg or two doses of 2 mg, was administrated to *Pvalb^FlpO^;Tcerg1l^CreER^;Ai65* mice of 1-2 month age. Tamoxifen solution was prepared by dissolving Tamoxifen (Sigma-Aldrich, T5648**)** in corn oil (Sigma-Aldrich) at 10-20 mg/ml concentration with agitation or sonication. Tamoxifen solution was either stored at RT and used within one week of preparation or stored long-term at −80°C and warmed up prior to injection. Tamoxifen solution was administrated to mice through oral gavage.

For induction of Kir2.1 expression, Tamoxifen was administered through intragastric or subcutaneous injection to *Fezf2^2A-CreER^;Rosa26^Kir2.1-mCehrry^* mice at P1. No Tamoxifen-induced pup mortality was observed. Due to the inherent variability of neonatal injection, successful induction was confirmed in each experimental animal by verifying the expression of *Kcnj2* and *mCherry* mRNAs using RNAscope.

**RNAscope *in situ* hybridization**

For fixed-frozen brain tissue, mice were deeply anesthetized with sodium pentobarbital (Euthasol) via intraperitoneal injection and transcardially perfused with 1X PBS followed by 4% paraformaldehyde (PFA). Brains were then dissected and post-fixed in 4% PFA overnight at 4°C. PFA-fixed brain samples were cryopreserved in 30% (w/v) sucrose and sectioned into 20 µm coronal slices using a sliding microtome (Leica). Brain slices were preserved in Storage Buffer, comprising 28% (w/v) sucrose, 30% (v/v) ethylene glycol in 0.1 M sodium phosphate buffer, at -80 °C until further processing. For fresh-frozen brain tissue, flash-frozen brain samples were sectioned on a cryostat (Leica) into 19-20 µm coronal slices. mRNA transcripts were detected using the RNAscope Multiplex Fluorescent V2 Assay Kit (ACDBio, 323100), following manufacturer’s protocol for either tissue type with following modification. Tissue was digested using Protease III for 30 min. at room temperature. The RNAscope catalogue probes used included *Lhx6* (#422791), *Gad2* (#439371), *Sst* (#404631), *Pvalb* (#421931), *Crhr2* (#413201), *Nmbr* (#406461), *Calb2* (#313641), *Hpse* (#412251), Kncj2 (#476261), mCherry (#431201).

**Image acquisition**

Images of RNAscope *in situ* hybridization experiments were collected using a tiling scope (Zeiss Axio Imager A1) with a 10X objective. Images of transgenic mouse line labeling were either collected using a whole slide scanning microscope with a 10X objective (Olympus VS120 slide scanners), or acquired with an upright confocal microscope (Zeiss LSM 800) with a 10X objective (Plan-Apochromat 10x/0.45 M27) or a 20X objective (Plan-Apochromat 20x/0.8 M27) to better appreciate the cellular morphology. Images of viral genetic labeling of SST-Hpse and SST-Crhr2 were imaged using another confocal microscope (Leica Stellaris) with a 10X objective (HC PL APO 10x/0.40 DRY). Stitching of image tiles were mostly performed using acquisition software, except for some used Stitching plugin in FiJi^1^.
